# The TONSL-MMS22L complex and FANCM form an interdependent complex on chromatin to counter replication stress

**DOI:** 10.1101/2025.03.07.642025

**Authors:** Haixia Zhou, Jiaoyan Yan, Xinlei Cao, Jingfei Zhan, Chen Ling, Yunhui Luo, Zhuang Ma, Yinyying Sun, Peining Song, Weiwei Liu, Lizhu Wang, Jiahuan Li, Althaf Shaik, Marina Bellani, Linqian Wang, Yun Xie, Jing Zhang, Xiangyang Xue, Xian Shen, Michael M. Seidman, Weidong Wang, Zhijiang Yan

**Affiliations:** Oncology Discipline Group, The Key Laboratory of Pediatric Hematology and oncology Diseases of Wenzhou, The Second Affiliated Hospital and Yuying Children’s Hospital of Wenzhou Medical University, China; Institute of DNA Repair Diseases, School of Basic Medical Sciences, Wenzhou Medical University, China; Laboratory of Genetics and Genomics, National Institute on Aging, National Institutes of Health, USA; Laboratory of Molecular Biology & Immunology, National Institute on Aging, National Institutes of Health, USA; Zhejiang Key Laboratory of Intelligent Cancer Biomarker Discovery and Translation, The First Affiliated Hospital of Wenzhou Medical University, Wenzhou, China; Tongji University School of Medicine, China

**Keywords:** FANCM, TONSL-MMS22L, replication stress, Fanconi Anemia

## Abstract

FANCM is branchpoint DNA translocase essential for cellular response to replication stress. Here, we show that replication stress stimulates FANCM and the TONSL-MMS22L heterodimer bound to histones H3-H4 to form an interdependent complex on chromatin. TONSL-MMS22L recruits FANCM and Fanconi anemia (FA) core complex to stalled and collapsed forks, maintains FANCM on replication-stressed chromatin, promotes FANCD2 monoubiquitination, facilitates both repair and replication traverse of DNA interstrand crosslinks (ICLs), and suppresses sister chromatid exchanges, through its interactions with FANCM and H3-H4. Reciprocally, both DNA translocase activity and phosphorylation of FANCM facilitate recruitment of TONSL-MMS22L and RAD51 to perturbed forks. Moreover, TONSL-MMS22L and FANCM function together to promote activation of the FA pathway, ICL repair, homologous recombination and replication traverse. Cancer patients with tumors with wildtype FANCM and low expression of TONSL-MMS22L have a more favorable prognosis than those with high expression. Thus, FANCM-TONSL-MMS22L acts coordinately as a complex on chromatin that resolves replication stress, and this complex may present a therapeutic target for wildtype FANCM-linked cancer.

## Introduction

Accurate and complete DNA replication is essential for genome integrity. The major challenges to the replication progression are fork stalling and collapsing, both of which are collectively referred to replication stress^1,2^. In vertebrates, one of the crucial players for resolving this stress is FANCM, which is a branchpoint DNA translocase and recruits distinct proteins involved in DNA damage response, repair and replication to coordinate cellular response to replication stress^3-7^. The human FANCM was initially identified as a member of the Fanconi anemia (FA) core complex^8^. FA is a rare hereditary bone marrow failure and cancer predisposition disease, caused by mutations in any of 22 FANC genes^9^. These FANC proteins facilitate a DNA damage response pathway called the FA pathway, which is critical to repair DNA interstrand crosslinks (ICLs) that covalently link two complementary strands of DNA and thus absolutely block replication^10,11^.

When replication forks encounter with ICLs, FANCM plays at least two essential roles. First, FANCM, along with its obligate partners (MHF and FAAP24), recognizes ICLs through its DNA binding activity^12,13^. Once bound to ICLs, FANCM recruits the FA core complex to bidirectional forks colliding with the same ICLs, consequently catalyzing monoubiquitination of FANCD2 and FANCI by UBE2T (FANCT)-FANCL to activate the FA pathway^11^. Second, FANCM, which is independent of the FA core complex, promotes replication machinery to traverse the unrepaired ICLs that stall single forks. This process, called replication traverse pathway, depends on the DNA translocase activity of FANCM, by which the replication is allowed to bypass ICLs and thus enables DNA synthesis to be completed. The ICLs are then removed by post-replication repair mechanisms^14^. Several FANCM-binding partners, including MHF, PCNA and BLM complex, work together with FANCM to facilitate traverse^14-16^. Beyond ICLs, FANCM participates in the other processes that eliminate replication stress. For example, promoting repair of collapsed forks provoked by topoisomerase I inhibitor camptothecin (CPT) and stalled forks induced by hydroxyurea (HU) or aphidicolin (APH)^12,17-19^, facilitating homologous recombination (HR) repair at ICL-independent double strand breaks (DSBs)^20^, suppressing sister chromatid exchanges^21,22^, and attenuating alternative lengthening of telomeres (ALT)^23-25^. The functions exerted by FANCM are largely relied on its two intrinsic determinants. One is the ATP-dependent branchpoint DNA translocase activity harbored by its N-terminal helicase domain, which promotes migration of branched DNA molecules, fork reversal, ATR activation and replication traverse of ICLs^6,7,26-28^. The other is its phosphorylation catalyzed by ATR, which facilitates cellular resistance to replication stress, activation of the FA pathway, FANCM recruitment to laser-induced ICLs, and replication traverse of ICLs^29,30^.

FANCM has been shown to bind chromatin regardless replication stress^12,31^. Despite the requirement of both MHF and FAAP24 for efficient association of FANCM with chromatin^12,31^, how FANCM controls both DNA repair and replication traverse pathways in the context of chromatin remains largely unknown. One possibility is due to a lack of prior discovery of chromatin structure-derived proteins and/or their bound proteins that interact with FANCM. In this study, we develop an immunopurification-based proteomic protocol to unbiasedly identify two novel FANCM cofactors, TONSL and MMS22L, from native chromatin of human cells under replication stress. These two proteins constitute an obligate heterodimer, called the TONSL-MMS22L complex, that is essential for HR repair by recruiting RAD51^32-36^. TONSL-MMS22L binds both soluble histones H3-H4 dimer that are unincorporated into nucleosomes^32,37^, and nucleosomal H3-H4 dimer in chromatin during replication^38^. Bi-allelic hypomorphic mutations in TONSL cause SPONASTRIME dysplasia and increase spontaneous chromosomal aberrations^39,40^. We provide evidence that TONSL-MMS22L inactivation disrupts FANCM retention on chromatin and impairs FANCM-mediated multiple DNA repair and replication traverse pathways upon replication stress. Mutually, FANCM loss impairs TONSL-MMS22L-RAD51 accumulation at perturbed forks and diminishes TONSL-MMS22L-mediated HR repair at DSBs. Furthermore, we present clinical evidence that TONSL-MMS22L low expression specifically depends on proficient FANCM, but not mutant FANCM, for a better prognosis of cancer patients.

## Results

### Replication stress stimulates the FANCM-TONSL-MMS22L complex on chromatin

We sought to identify novel FANCM-interacting proteins from chromatin under replication stress. However, the biochemical purification of the FANCM complex directly from chromatin is unfeasible, because of the insolubility of proteins due to their being packed into highly-ordered and compacted chromatin^41^. To solve this difficulty, we first isolated native chromatin of HeLa cells treated with hydroxyurea (HU) that induces replication stress by depleting nucleotide pools^42^, and then used microcococcal nuclease, followed by DNase I treatment, to release proteins from chromatin (Figure 1A). Since the yields of extracted proteins were too low to conduct an immunopurification-based mass spectrometry analysis, we employed gel filtration to concentrate the FANCM complex (Figure 1A). The purified FANCM complex from the pooled fractions containing FANCM (Figure 1A) with a FANCM antibody was subject to mass spectrometry. Unexpectedly, we identified TONSL and MMS22L in the FANCM complex (Figure 1B) and confirmed by immunoprecipitation (IP)-coupled western blotting with both FANCM and MHF1antibodies, but not pre-immune serum (Figure 1C). Moreover, both mass spectrometry and immunoblotting showed the presences of TONSL and MMS22L in the FANCM complex purified from HU-treated HEK293 cells stably expressing Flag-tagged FANCM with a Flag antibody (Figures S1A and S1B). In addition, the gel-filtrated profiles of TONSL and MMS22L were perfectly coincidental with that of FANCM (Figure 1A), providing extra evidence that TONSL and MMS22L are complexed with FANCM.

**Figure 1.**
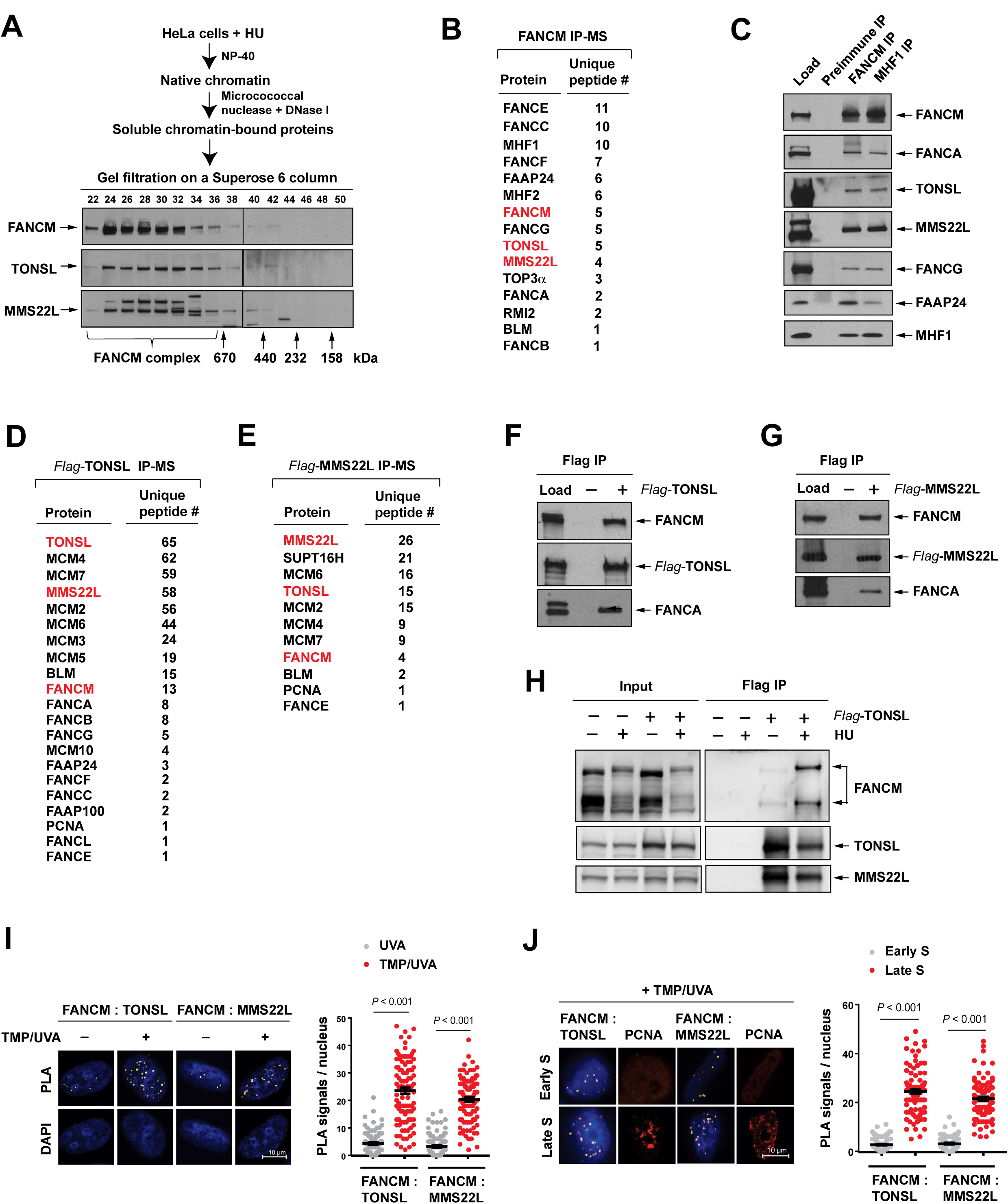
Replication stress stimulates the FANCM-TONSL-MMS22L complex. **(A)** A diagram depicting the isolation of soluble chromatin-bound proteins from HU-treated HeLa cells (*Top*), and immunoblotting showing the gel-fractionated profiles of FANCM, TONSL and MMS22L on a Superose 6 column (*Botton*). **(B)** Mass spectrometry showing the identified proteins and their corresponding number of unique peptides in the FANCM complex. **(C)** Immunoblotting showing the presences of TONSL, MMS22L and several known components of the FA core complex in the precipitates isolated with both FANCM and MHF1 antibodies, respectively. The IP with a preimmune serum was used as a control. **(D** and **E)** As described in (B), except the TONSL precipitate in (D) and the MMS22L precipitate in (E). **(F, G** and **H)** As described in (C), except a Flag antibody was used for IP. Cells without expression of any Flag-tagged protein were used as controls. **(I)** *Left panel:* Representative images of the PLA between FANCM and TONSL or MMS22L in HeLa cells with the indicated treatments. *Right panel:* A graph showing the frequencies of PLA signals from three biological replicates. Data are mean ± SEM. Number of nuclei: FANCM: TONSL with UVA = 105, TMP/UVA = 102; FANCM: MMS22L with UVA = 113, TMP/UVA = 107. **(J)** As described in (I), except the early and late phase cells were used. Number of nuclei: FANCM: TONSL in early S phase = 110, late S = 96; FANCM: MMS22L in early S phase = 94, late S = 99. (**See also Figure S1**)

To reciprocally validate the FANCM-TONSL-MMS22L association, we applied benzonase digestion to release proteins from native chromatin of HU-treated HeLa cells stably expressing Flag-TONSL or Flag-MMS22L. Both extracts were subjected to IP with a Flag antibody. Unexpectedly again, mass spectrometry revealed that both TONSL and MMS22L precipitates contained FANCM, BLM, PCNA and multiple components of the FA core complex (Figures 1D and 1E). The appears of FANCM and FANCA in both the precipitates were verified by immunoblotting (Figures 1F and 1G). These data demonstrate that the FANCM-TONSL-MMS22L complex is indeed integral to the FA core complex.

After establishing the association between FANCM and TONSL-MMS22L following HU treatment, we examined whether this complex was influenced by replication stress. First, we performed IPs of Flag-tagged TONSL from both HU-treated and -untreated cells with a Flag antibody. The following immunoblotting revealed a small amount of FANCM in the TONSL precipitate isolated from untreated cells and a considerable amount from treated cells, but not from both untreated and treated control cells (Figure 1H). Second, we employed the proximity ligation assays (PLAs), which are a powerful tool to detect *in situ* protein-protein interactions in cells^43^. Before conducting PLA, HeLa cells were treated with trimethylpsoralen (TMP), followed by long wave ultraviolet light (UVA) irradiation, which can randomly produce ICLs between two complementary strands of DNA that induces severe replication stress^44^. Using UVA irradiation alone as a control because it does not induce replication stress^45^, we observed that the PLA signals that reflect the FANCM-TONSL-MMS22L interactions were low in UVA-treated cells, but sharply increased by ∼ 20-fold in TMP/UVA-treated cells (Figure 1I), in agreement with the IPs (Figure 1H). Together, the data indicate that the FANCM-TONSL-MMS22L complex is constitutive, albeit low, while considerably enhanced when cells suffer from replication stress.

### TONSL-MMS22L specifically associates with FANCM-bound stressed replisomes

FANCM is predominantly bound to the proximal replisomes at ICL-stalled forks in heterochromatin in late S phase of the cell cycle^46^. We performed PLAs with TMP/UVA-treated early and late S phase cells sorted by flow cytometry. Using PCNA as a marker of recruitment to damaged DNA prominently in late S phase^47^, the PLA signals between FANCM and TONSL-MMS22L were increased by ∼ 20-fold in late S phase compared to those in early S phase (Figure 1J), whereas much less PLA signals between GFP-DONSON and TONSL-MMS22L were detected in both early and late S phases (Figure S1C). DONSON is another genome replication factor required for cellular response to replication stress and was included as a control^46,48^. From these data, we therefore conclude that TONSL-MMS22L retains specifically on FANCM-bound stressed replisomes in heterochromatin in late S phase.

### Both TONSL and MMS22L heterozygous knockout cells are hypersensitive to replication stress

We attempted to create TONSL- or MMS22L-null HeLa cells by CRISPR/Cas9 editing. Albeit successful in generating homozygous knockouts of FANCM, FANCD2 or FANCA (Figures S2A - S2C), we failed to obtain any TONSL- or MMS22L-null clones with three distinct gRNAs. Indeed, biallelic inactivation of TONSL (*TONSL^-/-^*) in both mice and zebrafishes by CRISPR resulted in early embryonic lethality^39^. These combined data indicate that both TONSL and MMS22L are most likely essential for cell viability among species.

Knockdown of TONSL or MMS22L rendered cells hypersensitive to CPT^32-35^. Thus, we picked up the CRISPR clone with a low level of TONSL (clone TG1-5) (Figure S2D) or MMS22L (clone MG2-2) (Figure S2F) and measured their sensitivity to CPT. Surprisingly, both TG1-5 and MG2-2 clones exhibited hypersensitivity to CPT, equivalent to TONSL- or MMS22L-knockdown cells (Figures S2H and S2I). Importantly, this sensitivity was corrected largely by re-expression of TONSL and partially of MMS22L, respectively (Figures S2H, S2I, 2B and S4A), ruling out the possibility of an off-target effect on CPT sensitivity by CRISPR editing. Sequencing reveals that TG1-5 is a heterozygous knockout, marked as *TONSL^+/-^* (Figure S2E); MG2-2 is a compound heterozygote, marked as *MMS22L*^Δ/-^ (Figure S2G). We, therefore, established both *TONSL^+/-^* and *MMS22L*^Δ/-^ as hypomorphic mutants for study thereafter.

### TONSL-MMS22L promotes recruitment of FANCM and FA core complex to perturbed replication forks

TONSL contains several conserved protein-protein interacting motifs and thus acts as a scaffold for anchoring multiple proteins, such as the N-terminal eight tetratricopeptide repeats (TPR) that associate with multiple MCM proteins and bind histone H3.1 variant on newly-replicated chromatin, the middle ankyrin repeat domain (ARD) that binds both nonnucleosomal histones H3-H4 dimer and H3-H4 dimer on post-replicative chromatin by recognizing unmethylated histone H4 at lysine 20 (H4K20me0), and the C-terminal seven leucine-rich repeats (LRR) that interact with MMS22L (Figure 2A)^32,34,38,49,50^. This prompted us to reason that TONSL might mediate the interaction between TONSL-MMS22L and FANCM. Indeed, we mapped the FANCM-interaction domain within TONSL to a minimum region covering residues Glu661 to Gly670, because deletion of this region completely abolished the interaction between TONSL and FANCM, as detected by IP-western blotting (Figure 2A). This region (aa661-670) is ranged out of the highly conserved residues in ARD of TONSL that are required for H4-tail binding^38^. Thus, these data reveal that TONSL links TONSL-MMS22L to FANCM via its region spanning aa 661 to 670.

**Figure 2.**
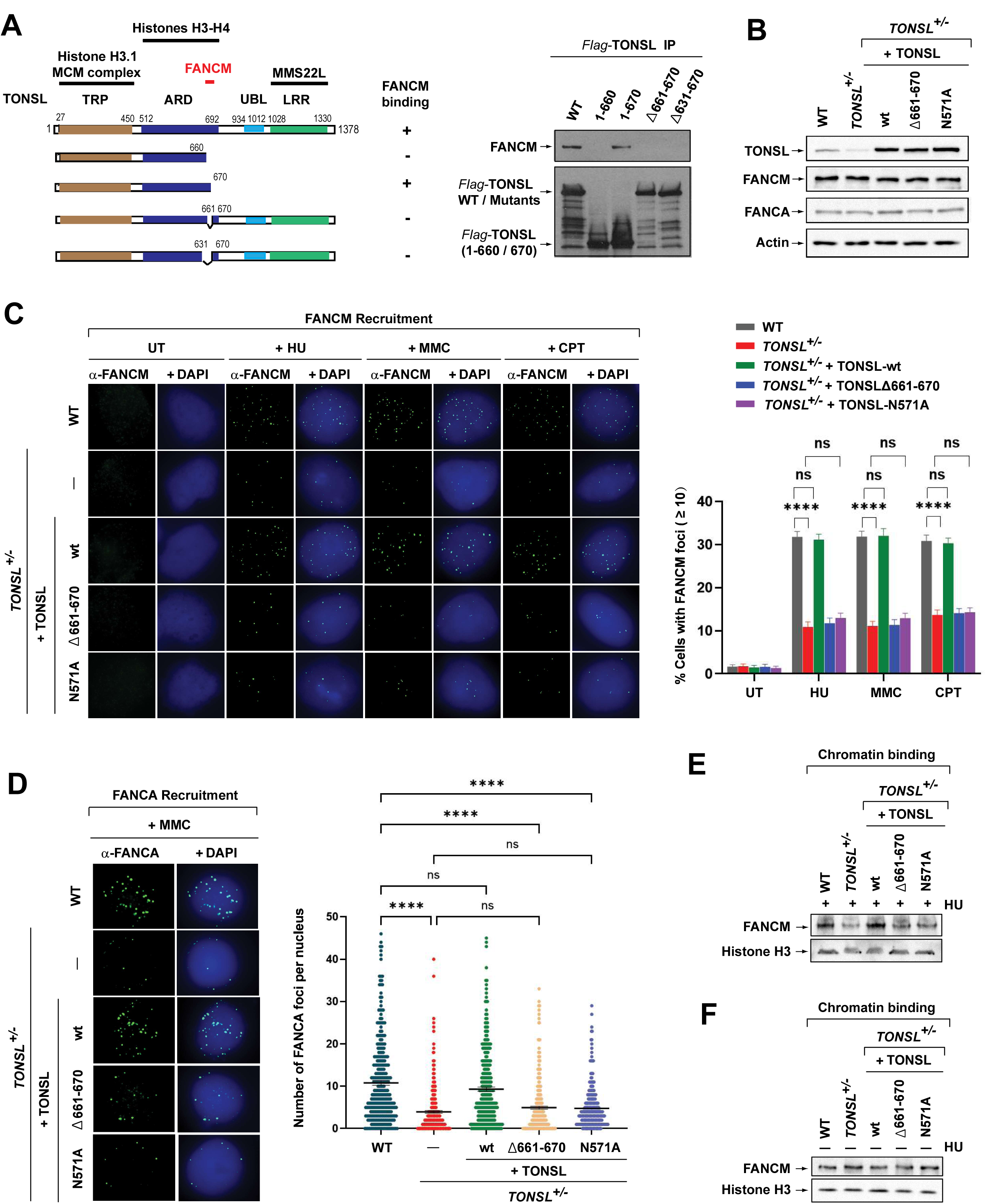
TONSL-MMS22L recruits FANCM and the FA core complex to stalled and collapsed forks, and maintains FANCM on replication-stressed chromatin through its interaction with FANCM and histones H3-H4. **(A)** A diagram showing the mapping of the FANCM-interacting domain within TONSL. *Left panels* show Flag-tagged wildtype and various deletion mutants of TONSL used in IP-Western blotting as indicated in *Right panels*. *Middle panels* show the interactions or non-interactions between various TONSL constructs and FANCM. *Right panels* showing the IP-Western blotting to map the FANCM-interacting domain within TONSL. Flag-tagged TONSL wildtype (WT) and indicated deletion mutants were transiently expressed in HeLa cells, respectively. Immunoprecipitation was carried out using anti-Flag M2 agarose beads. The presence of various TONSL fragments and FANCM were detected by Western blotting. A diagram summarizing the FANCM-TONSL interaction data is shown in the *Middle panels*. **(B)** Immunoblotting shows the levels of TONSL, FANCM and FANCA in whole cell lysates from HeLa wildtype cells (WT), *TONSL^+/-^*, and *TONSL^+/-^* cells complemented with TONSL wildtype (wt), TONSLΔ661-670 or TONSL-N571A mutant. Actin was included as a loading control. **(C)** Immunofluorescence images (*Left panel*s) and a quantification graph (*Right panels*) showing FANCM nuclear foci in various cells described in (B) after treatment with HU (2 mM) for 24 h, MMC (60 ng/ml) for 18 h or CPT (1.5 μM) for 24 h. Data are mean ± SEM from three independent experiments. **** represents *P* < 0.0001. “ns” represents nonsignificant difference. **(D)** As described in (C), except a FANCA antibody and MMC were used. **(E)** Immunoblotting shows the level of FANCM in chromatin fraction of HU-treated various cells described in (B). Histone H3 was included as a loading control. **(F)** As described in (E), except without HU treatment. **(See also Figures S2, S3 and S4)**

We have previously shown that the chicken endogenous FANCM was recruited as nuclear foci to stalled forks in avian DT40 cells^16^. However, such focus formation in human cells remains unreported. This might be due to a lack of an antibody against human FANCM for immunofluorescence (IF). We thus engineered this antibody. Equal to chicken FANCM, we demonstrated the recruitment of human endogenous FANCM to stalled forks induced by HU or mitomycin C (MMC) and to collapsed forks provoked by CPT (Figures S3A-S3C). This recruitment requires both the DNA translocase activity and phosphorylation of FANCM (Figures S3A-S3C).

After establishing FANCM recruitment in human cells, we assessed whether the FANCM-TONSL-MMS22L interaction is needed for this recruitment. Intriguingly, following HU, MMC or CPT treatment, both *TONSL^+/-^* and *MMS22L*^Δ/-^ cells showed all marked reductions in the population of cells with more than 10 of FANCM foci compared to WT cells (∼ 31% *vs* 12%) (Figures 2B, 2C and S4A-C). Importantly, these declines were all fully corrected by reintroduction of wildtype TONSL or MMS22L, but completely not of the TONSLΔ661-670 mutant that disrupts the TONSL interaction with FANCM (Figures 2B, 2C and S4A-C). Given that FANCM acts upstream the FA core complex^11^, we next examined whether the FANCM-TONSL-MMS22L complex contributes to recruitment of the FA core complex to forks stalling by ICLs. By measuring the FANCA nuclear foci that serve as a characteristic marker for the focus formation of the FA core complex in response to DNA damage^51^, both *TONSL^+/-^* and *MMS22L*^Δ/-^ cells revealed that MMC-induced FANCA foci were reduced by ∼2.7 and ∼ 2.1-fold, respectively; and restored by reintroduction of wildtype TONSL or MMS22L (Figures 2D and S4D), but completely not of the TONSLw661-670 mutant (Figure 2D). Of note, these alterations between reduction and restoration in both FANCM and FANCA foci were not duo to the stability of FANCM and FANCA, because the levels of both proteins were comparable among these cell types (Figures 2B and S4A). Therefore, the interaction between FANCM and TONSL-MMS22L is indispensable for efficient recruitment of FANCM and the FA core complex to stalled and collapsed forks.

### The histone-binding activity of TONSL-MMS22L is required for recruitment of FANCM and FA core complex

During DNA replication, TONSL-MMS22L binds histones H3-H4 incorporated into new nucleosomes through TONSL recognizing H4K20me0 in newly-replicated chromatin via its ankyrin repeat domain (ARD)^38^. Substitution of Asparagine (N)571 with Alaine (A) in ARD abolished TONSL binding to H4K20me0, resulting in disruption of TONSL-MMS22L association with chromatin and accumulation at collapsed forks as well as in rendering cells hypersensitive to CPT^38^. We wished to know whether this histone-binding activity is important for recruitment of FANCM and the FA core complex to stressed forks. Thus, we ectopically expressed TONSL-N571A mutant that inactivates TONSL-MMS22L-chromatin association in *TONSL^+/-^* cells (Figure 2B). Unlike TONSL wildtype, re-expression of this mutant completely failed to restore the reductions in both FANCM and FANCA foci seen in *TONSL^+/-^* cells after exposure to HU, MMC or CPT (Figures 2B, 2C and 2D), indicating that TONSL-MMS22L requires its histone-binding activity to recruit FANCM and the FA core complex to stalled and collapsed forks.

### FANCM retention on replication-stressed chromatin depends on TONSL-MMS22L and its histone-binding activity

Next, we carried out subcellular fractionation under normal and replication-stressed conditions to examine whether TONSL-MMS22L is necessary for FANCM retention on chromatin. Immunoblotting showed that a profoundly reduced level of FANCM in chromatin fraction was detected in *TONSL^+/-^* cells after HU treatment, compared to that in WT cells (Figure 2E). Notably, reintroduction of TONSL wildtype, but not both TONSLΔ661-670 and TONSL-N571A mutants, restored this reduction to level detected in WT cells (Figure 2E), consistent with the FANCM recruitment (Figure 2C). These data demonstrate that TONSL-MMS22L requires its interaction with FANCM or histone-binding activity to promote FANCM binding to chromatin upon replication stress. In contrast, the level of FANCM in chromatin fractions was comparable among these TONSL-altered cell types in the absence of HU treatment (Figure 2F), indicating that TONSL-MMS22L is dispensable for FANCM association with chromatin under normal replication.

### TONSL-MMS22L is required for normal activation of the FA pathway, ICL repair and full suppression of SCEs

Our findings that TONSL-MMS22L is an integral component of the FA core complex (Figures 1B-1G) and required for recruitment of FANCM and the FA core complex to stalled forks (Figures 2C, 2D and S4B-S4D) prompted us to study if this complex is needed for activation of the FA pathway. First, we found that depletion of TONSL or MMS22L by two different siRNAs led to a reduced level of monoubiquitinated FANCD2 following exposure to MMC or HU (Figures 3A and S5A). Second, a reduction in FANCD2 monoubiquitination in both *TONSL^+/-^* and *MMS22L*^Δ/-^ cells was detected after HU treatment (Figures 3B and S5B). Third, the formation of FANCD2 nuclear foci, which is another obligate marker for activation of the FA pathway^52^, was markedly declined in both *TONSL^+/-^* and *MMS22L*^Δ/-^ cells after treatment with MMC or HU (Figures 3C and S5C). Strikingly, these defects were corrected by re-expression of wildtype TONSL (Figures 3B and 3C) or MMS22L (Figures S5B and S5C), but completely not of both TONSLΔ661-670 and TONSL-N571A mutants (Figures 3B and 3C). Together, the results demonstrate that TONSL-MMS22L, resembling other components of the FA core complex^11^, is required for normal function of the FA pathway, and that this requirement depends on its interaction with FANCM or histone-binding activity.

**Figure 3.**
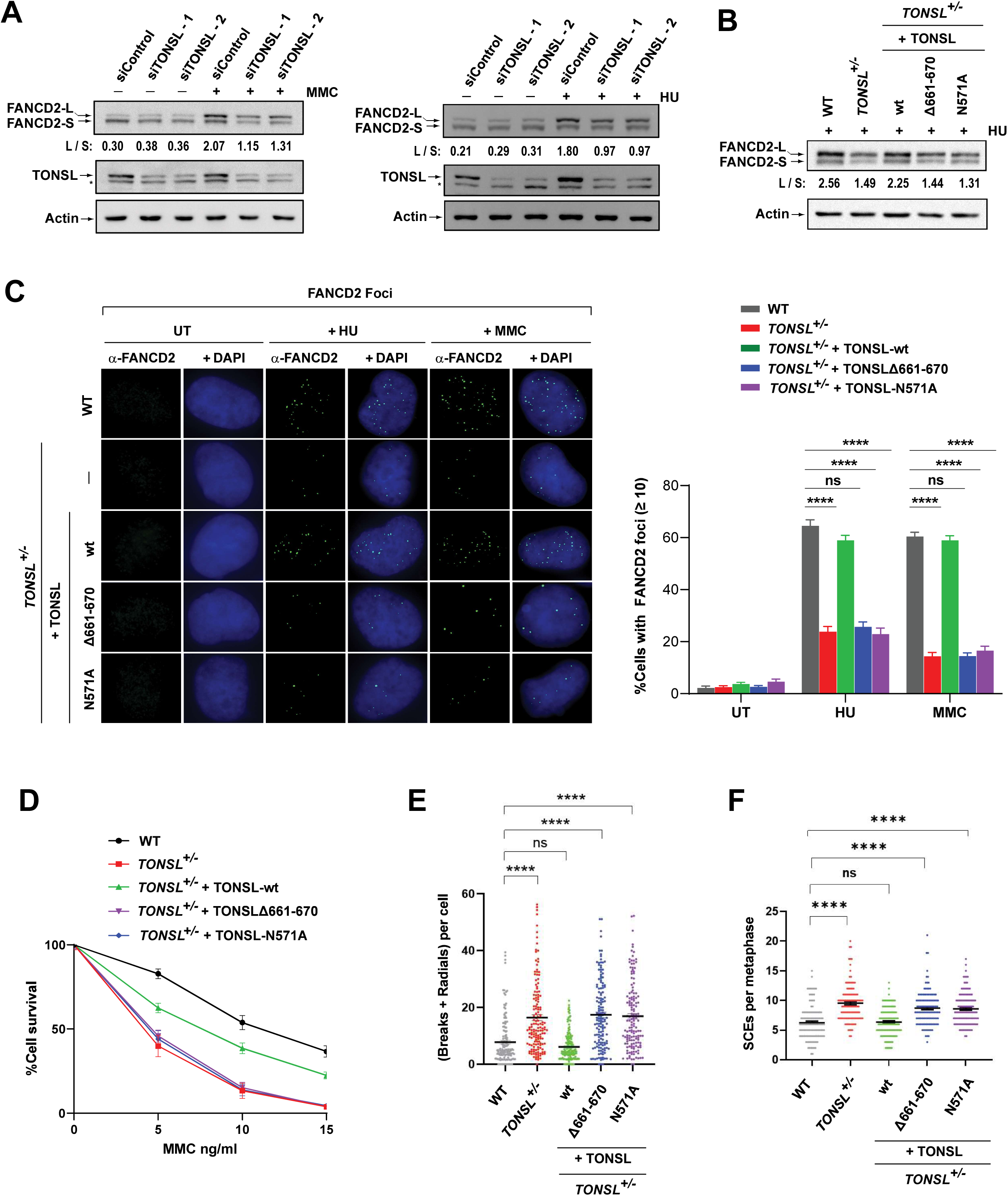
TONSL-MMS22L promotes activation of the FA pathway, ICL repair and SCE suppression through its interaction with FANCM and histones H3-H4. **(A)** Immunoblotting shows that HeLa cells depleted of TONSL have a reduced level of monoubiquitinated FANCD2 in the presence of MMC (60 ng/ml) for 16 h (*Left panel*) or HU (2 mM) for 16 h (*Right panel*). “L” (long) and “S” (short) represent ubiquitinated and non-ubiquitinated forms, respectively. The ratio between long and short forms was obtained using Image J software and shown below the blots. Actin was used as a loading control. **(B)** As described in (A), except HeLa wildtype cells (WT), *TONSL^+/-^* cells and *TONSL^+/-^* cells complemented with wildtype (wt) or mutant TONSL as described in Figure 2B. **(C)** Immunofluorescence images (*Left panels*) and a quantification graph (*Right panels*) showing FANCD2 nuclear foci in various cells described in Figure 2B after treatment with HU (2 mM) for 16 h or MMC (60 ng/ml) for 16 h. Data are means ± SEM from three independent experiments. **** represents *P* < 0.0001. “ns” represents nonsignificant difference. **(D)** Clonogenic survival assays of indicated cells following MMC treatment at the indicated concentrations. Data are mean ± SEM from three independent experiments. Each experiment has triplicate cultures. **(E)** A graph showing levels of MMC-induced chromosomal breaks and radial chromosomes of various cells as indicated. Data are mean ± SEM from three independent experiments. About 50 metaphase cells were counted per sample in each experiment. **** represents *P* < 0.0001. “ns” represents nonsignificant difference. **(F)** As described in (E), excepted the levels of spontaneous SCEs. **(See also Figure S5)**

We next found that both *TONSL^+/-^* and *MMS22L*^Δ/-^ cells exhibited hypersensitivity to both MMC and cisplatin that induce ICLs (Figures 3D, S5D and S5E), and profoundly increased frequencies of chromosomal breaks and radials by ∼ 2 and 1.7-fold, respectively, compared to WT cells after MMC treatment (Figures 3E, S5F and S5G). Moreover, both *TONSL^+/-^* and *MMS22L*^Δ/-^ cells revealed a higher level of SCEs (∼ 1.5-fold) than WT cells (Figures 3F, S5H and S5I). Likewise, re-expression of wildtype TONSL or MMS22L, but not both TONSLΔ661-670 and TONSL-N571A mutants, restored these defective phenotypes (Figures 3D-3F, S5D - S5I), indicating that TONSL-MMS22L requires its interaction with FANCM or histone-binding activity to promote ICL repair and fully suppress SCEs.

### TONSL-MMS22L works together with FANCM and the FA core complex for FANCD2 monoubiquitination, cellular resistance to replication stress and SCE suppression

We performed genetic dissection to investigate how TONSL-MMS22L works with FANCM and the FA core complex. Strikingly, depletion of TONSL or MMS22L in *FANCM^-/-^* cells resulted in no additive impairments of FANCD2 monoubiquitination and CPT sensitivity compared to those in single gene-deficient cells (Figures 4A, 4C, S6A and S6C). Similarly, *TONSL^+/-^* or *MMS22L*^Δ/-^ cells depleted of FANCA displayed no further reductions in FANCD2 monoubiquitination and MMC sensitivity (Figures 4B, 4D, S6B and S6D). FANCM depletion in both *TONSL^+/-^* and *MMS22L*^Δ/-^ cells led to no additional impairments of MMC sensitivity and SCE frequencies (Figures 4E, 4F, S6E and S6F). Taken together, we conclude that TONSL-MMS22L works with FANCM and the FA core complex in the same pathway for activation of the FA pathway, cellular resistance to replication stress and SCE suppression.

**Figure 4.**
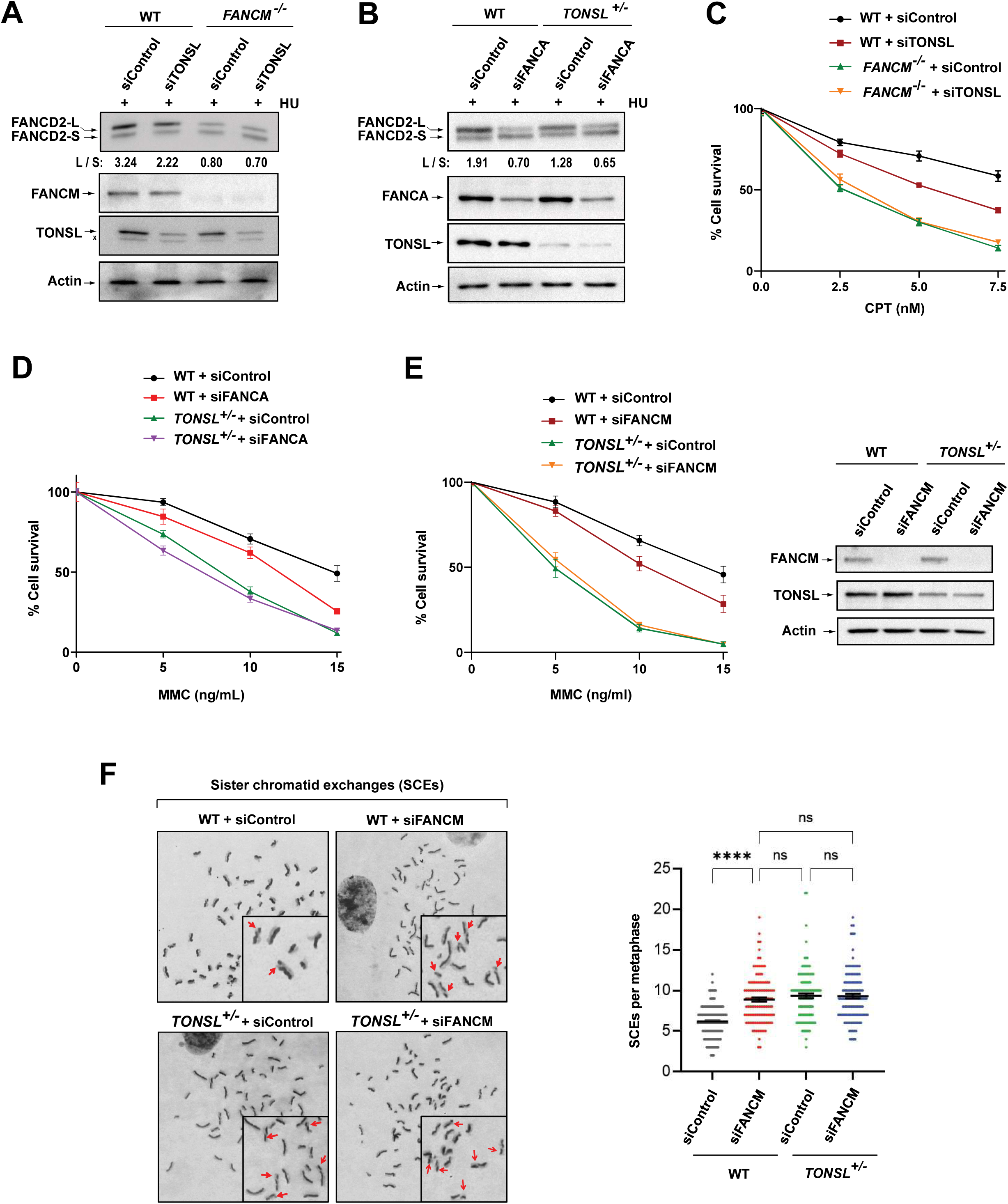
TONSL-MMS22L works together with FANCM and the FA core complex for FANCD2 monoubiquitination, cellular resistance to replication stress and SCE suppression. **(A)** Immunoblotting shows the levels of monoubiquitinated and unubiquitinated FANCD2, FANCM and TONSL in whole cell lysates from various cells as indicated. Cells were treated with HU (2 mM) for 18 h. Actin was used as a loading control. **(B)** As described in (A), except FANCA. **(C)** Clonogenic survival assays of indicated cells as following CPT treatment at the indicated concentrations. Data are mean ± SEM from three independent experiments. Each experiment has triplicate cultures. **(D)** As described in (C), except that MMC was used. **(E)** *Left panels:* As described in (C), except MMC was used. *Right panels:* Immunoblotting shows the levels of FANCM and TONSL in whole cell lysates from various cells as indicated. Actin was used as a loading control. **(F)** *Left panel:* Representative images showing SCEs (red arrows) in various cells as indicated. *Right panel:* A graph showing the spontaneous SCE levels of various cells as indicated. Data are mean ± SEM from three independent experiments. About 50 metaphase cells were counted per sample in each experiment. **** represents *P* < 0.0001. “ns” represents nonsignificant difference. **(See also Figure S6)**

### TONSL-MMS22L inactivation exposes its indispensability for replication traverse of ICLs

One of the defined functions of FANCM is required for replication machinery to traverse ICLs when the single forks encounter with these lesions^14^. To examine whether this traverse requires TONSL-MMS22L, we performed ICL-coupled DNA fiber assays as previously described^14^. Briefly, cells were treated with digoxigenin-labeled trimethylpsoralen (Dig-TMP), followed by UVA irradiation, to randomly create ICLs in DNA, and then sequentially pulsed with CIdU and IdU to label DNA replication tracts. The pattern of each labeled fork embedded with the labeled ICL was visualized by IF with stretched DNA fibers (Figure 5A). Interestingly, the traverse frequencies were markedly reduced in both *TONSL^+/-^* and *MMS22L*^Δ/-^ cells compared to those in WT cells (∼ 57% *vs* 34%) (Figures 5B and 5C). The reduction ratio was comparable to those in FANCM knockout HeLa, MEF or DT40 cells and in FANCM knockdown HeLa cells (∼ 40 - 50% of reduction)^15,16,30,53^. Importantly, re-expression of wildtype TONSL and MMS22L restored these declines to levels equal to those in WT cells, respectively (Figures 5B and 5C), whereas the traverse frequency in *TONSL^+/-^* cells expressed of either TONSLΔ661-670 or TONSL-N571A mutant remained the similar level to that in *TONSL^+/-^* cells (∼ 32% *vs* 34%) (Figure 5B). Thus, TONSL-MM22L, like FANCM, is required for promoting replication traverse of ICLs through its interaction with FANCM or its histone-binding activity.

**Figure 5.**
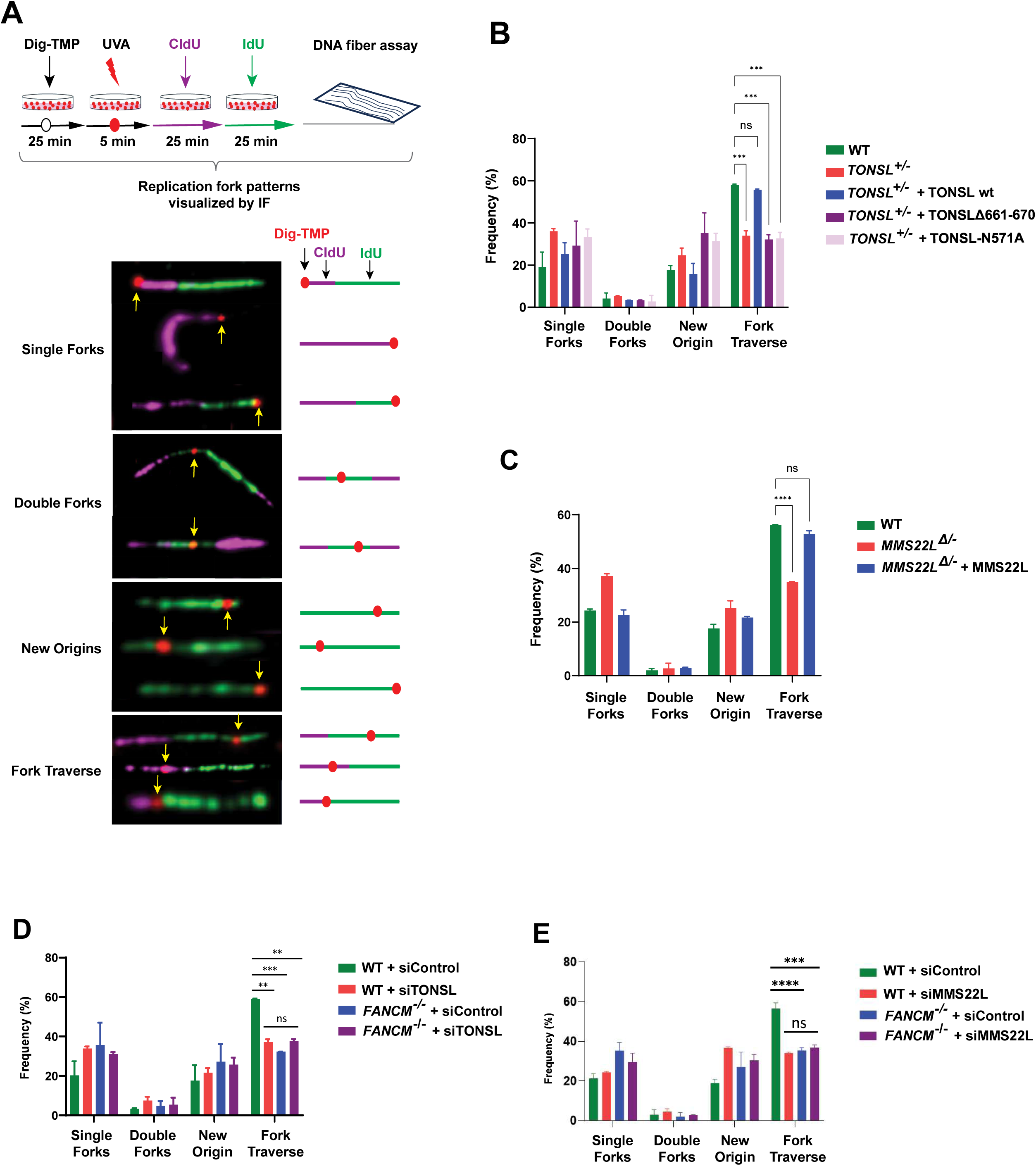
TONSL-MMS22L and FANCM act in the same pathway to promote replication traverse of ICLs. **(A)** (*Top panel*) A diagram showing the protocol of replication traverse assay. (*Botton panel*) Representative images showing the patterns of replication tracts in the vicinity of Dig-TMP-induced ICLs (marked with red dots) on DNA fibers. The sequentially purple-colored CIdU and green-colored IdU tracts define the directions of replication. **(B)** A graph showing the pattern distributions indicated in (A). Number of fibers with ICLs in indicated cells from two independent experiments: WT = 151, 137; *TONSL^+/-^* = 193, 146; *TONSL^+/-^* + TONSL-wt = 150, 148; *TONSL^+/-^* + TONSLΔ661-670 = 157, 144; *TONSL^+/-^* + TONSL-N571A =147, 147. Data are mean ± SEM. **(C)** As described in (B), except number of fibers: WT = 155, 158; *MMS22L*^Δ/-^ = 154, 150; *MMS22L*^Δ/-^ + MMS22L = 164, 150. **(D)** As described in (B), except number of fibers: WT + siControl = 138, 142; WT + siTONSL = 158, 146; *FANCM^-/-^* + siControl = 152, 167; *FANCM^-/-^* + siTONSL = 138, 151. **(E)** As described in (B), except number of fibers: WT + siControl = 162, 165; WT + siMMS22L = 166, 153; *FANCM^-/-^* + siControl = 170, 151; *FANCM^-/-^* + siMMS22L = 183, 169.

### TONSL-MMS22L and FANCM operate in the same pathway to promote ICL traverse

We next performed traverse assays with single and double gene-deficient cells simultaneously. Cells depleted of TONSL or MMS22L showed a marked reduction in traverse events compared to WT cells (siTONSL: ∼ 59% *vs* 37%; siMMS22L: ∼ 57% *vs* 34%) (Figures 5D and 5E), consistent with the findings in *TONSL^+/-^* and *MMS22L*^Δ/-^ cells (Figures 5B and 5C). Importantly, depletion of TONSL or MMS22L in *FANCM^-/-^* cells resulted in no further impairment of traverse compared to any of single gene-deficiency (Figures 5D and 5E), demonstrating that TONSL-MMS22L and FANCM are epistatic for ICL traverse.

### FANCM loss impairs recruitment of TONSL-MMS22L and RAD51 to perturbed forks

Upon replication stress, TONSL-MMS22L is recruited to stalled and collapsed forks, and DSBs to load RAD51 onto ssDNA as nuclear foci^34,36^. We asked whether FANCM is needed for this recruitment. Compared to WT cells, *FANCM^-/-^* cells showed a profound reduction in the population of cells with more than five of TONSL or MMS22L foci by ∼ 50% after treatment with MMC or CPT (Figures 6A and S7A). Strikingly, reintroduction of wildtype FANCM restored these declines to levels equivalent to those seen in WT (Figures 6A and S7A), whereas re-expression of either DNA translocase-inactivated mutant (K117R)^8^ or phosphorylation-abolished mutant (S1045A)^29^ of FANCM was completely incapable of correcting the reductions in both TONSL and MMS22L foci observed in *FANCM^-/-^* cells (Figures 6A and S7A). Similarly, compared to WT cells, the percentage of *FANCM^-/-^* cells with more than 10 of RAD51 foci was reduced by nearly four-fold (∼ 32% *vs* 8%), and fully rescued by re-expression of wild type FANCM, but completely not of either FANCM-K117R mutant or FANCM-S1045A mutant (Figure 6B). Taken together, the results indicate that FANCM is required for full recruitment of TONSL-MMS22L and RAD51 to perturbed forks, and that this recruitment is entirely manipulated by its DNA translocase activity or phosphorylation.

**Figure 6.**
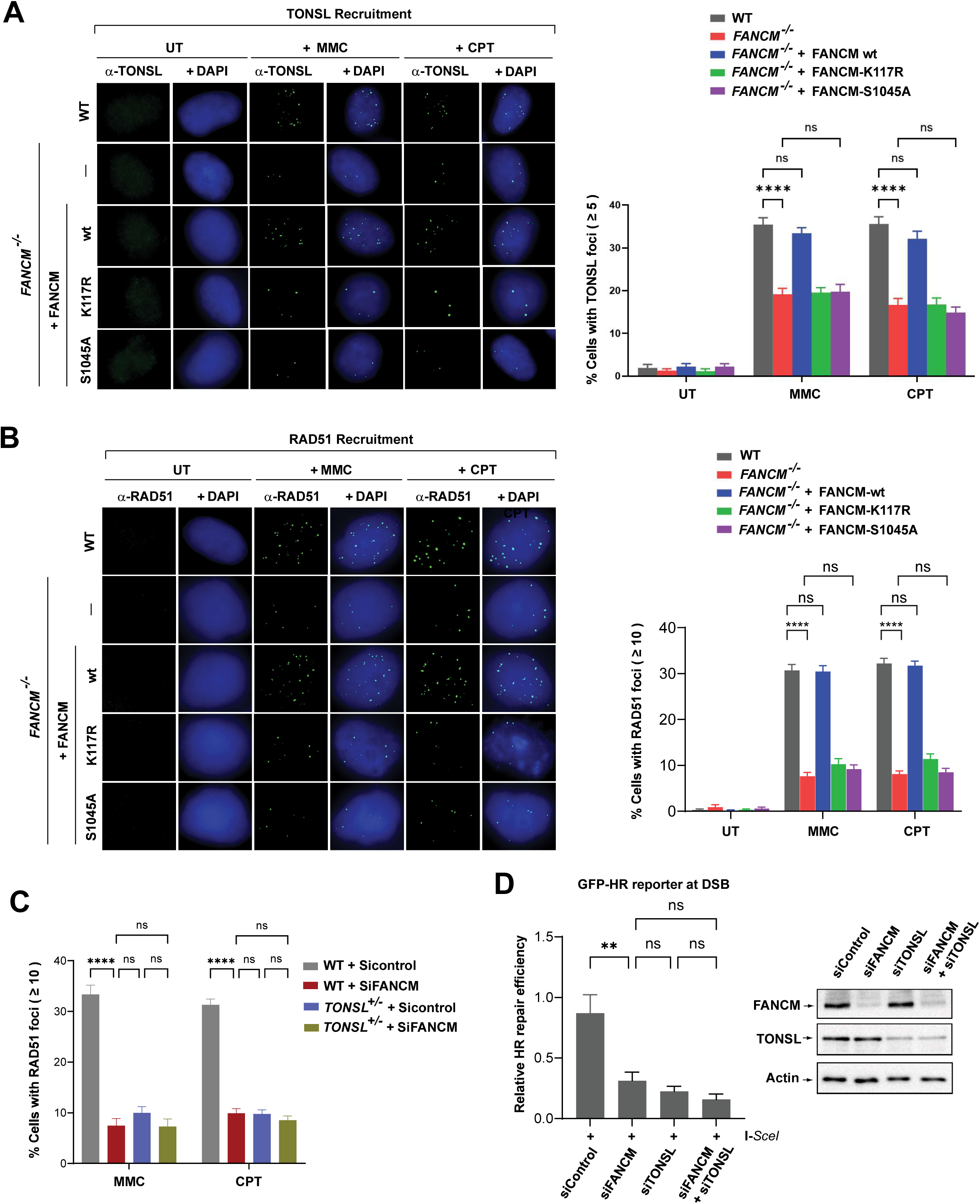
FANCM and TONSL-MMS22L function together to facilitate HR repair. **(A)** Representative images (*Left panel*) and a quantification graph (*Right panel*) showing TONSL nuclear foci in various cells as indicated after treatment with MMC (60 ng/ml) for 18 h or CPT (1.5 μM) for 24 h. Data are mean ± SEM from three independent experiments. **** represents *P* < 0.0001. “ns” represents nonsignificant difference. **(B)** As described in (A), except RAD51 nuclear foci and CPT (50 nM) treatment for 2 hr. **(C)** As described in (A, *Right panel*), except RAD51 nuclear foci. **(D)** *Left panels:* A quantitation graph showing relative repair efficiency derived from the positive GFP cells in U2OS-DR-GFP-reporter cells transfected with various siRNA oligos as indicated. Data are the mean with SEM from three independent experiments. ** represents *P* < 0.01. “ns” represents nonsignificant difference. *Right panels:* Immunoblotting shows the level of FANCM and TONSL in whole cell lysates of various cells as indicated. Actin was included as a loading control. **(See also Figure S7)**

### FANCM and TONSL-MMS22L act in the same pathway for RAD51 recruitment to perturbed forks

Since both FANCM and TONSL-MMS22L are necessary for RAD51 recruitment (Figure 6B)^34,36^, we wondered how they work genetically in this scenario. We depleted FANCM in WT, *TONSL^+/-^*or *MMS22L*^Δ/-^ cells by siRNA and examined RAD51 focus formation following treatment with MMC or CPT. In agreement with the findings in *FANCM^-/-^* cells (Figure 6B), FANCM depletion also resulted in a nearly four-fold reduction in the percentage of cells with more than 10 of RAD51 foci compared to siControl (Figures 6C and S7B). Likewise, approximately 8-10% of both *TONSL^+/-^*and *MMS22L*^Δ/-^ cells treated with siControl exhibited more than 10 of RAD51 foci per cell, while 32% of WT cells were positive (Figures 6C and S7B), consistent with the previous reports that knockdown of TONSL-MMS22L led to a decline of RAD51 foci^34^. Importantly, either *TONSL^+/-^* or *MMS22L*^Δ/-^ cells depleted of FANCM remained the similar level of reduction in RAD51 foci to that seen in any type of single gene-deficient cells (Figures 6C and S7B). Thus, TONSL-MMS22L and FANCM are epistatic for RAD51 loading onto ssDNA at DSB ends derived from forks stalling by MMC-induced ICLs or forks collapsing provoked by CPT.

### FANCM and TONSL-MMS22L function together to promote homologous recombination (HR) repair

Depletion of TONSL-MMS22L impaired RAD51 loading and consequently attenuated HR repair at DSBs, when replication forks suffer from stalling or collapsing^34^. We thus attempted to investigate whether FANCM plays a role in DSB repair by promoting HR, because loss of FANCM led to profound reductions in recruitment of both TONSL-MMS22L and RAD51 to stalled and collapsed forks. We depleted FANCM in U2OS cells carrying I-SceI-based GFP-HR reporter construct and performed cell-based DSB repair assays. Interestingly, depletion of FANCM resulted in a nearly two-fold reduction in HR efficiency compared to siControl (Figures 6D and S7C), in agreement with the recent note that FANCM along with FANCL and UBE2T were identified as HR promoting factors in human cells^20^. As expected, U2OS cells depleted of TONSL or MMS22L exhibited a marked reduction in HR repair activity (Figures 6D and S7C), consistent with the previous report^34^. Strikingly, double depletion of either both FANCM and TONSL or both FANCM and MMS22L resulted in no additive impairment of HR repair compared to depletion of single gene (Figures 6D and S7C). Given that FANCM promotes recruitment of both TONSL-MMS22L and RAD51 to forks stalling by MMC-induced ICLs and to collapsed forks-derived DSBs by CPT collision (Figures 6A, 6B and S7A), and that FANCM and TONSL-MMS22L work together to facilitate the RAD51 recruitment (Figures 6C and S7B), these data suggest that FANCM and TONSL-MMS22L are epistatic for HR repair at both ICL-dependent and -independent DSBs.

### Patients with low TONSL-MMS22L expression in tumors with wild type FANCM have a more favorable prognosis than those with high expression

Mutations in FANCM can pose a high risk for onset of breast cancer and other types of cancer in both human and mouse^3^. Our findings that TONSL-MMS22L interacts not only physically but also functionally with FANCM to relieve replication stress prompted us to ask whether TONSL-MMS22L has a pathological relevance to FANCM in clinic. To address this, we performed the survival analyses of the Cancer Genome Atlas (TCGA) patients. First, using the clinical and genomic alteration data of TCGA Pan-Cancer on 32 types of cancer, 310 samples of cancer patients harboring FANCM mutation who suffered from 21 types of cancer with or without therapeutic treatments were dug out. Accordingly, the same size of patient samples with wildtype FANCM along with the same genders, types of cancer and similar ages as those with mutated FANCM were randomly mined out. Then, using mRNA expression Z-scores of TCGA Pan-Cancer on 32 types of cancer, patients with wildtype or mutant FANCM were divided into the groups with low or high expression of either TONSL or MMS22L. Interestingly, in cancer patients with wildtype FANCM, the group with low expression of TONSL or MMS22L showed a longer survival ratio than the group with high expression (*P* < 0.05) (Figure 7A); whereas, in cancer patients with mutant FANCM, no correlation between TONSL-MMS22L expression and overall survival was detected (*P* > 0.05) (Figure 7B). To rule out the non-specificity for these analyses, we used a characteristic member of the FA core complex, FANCA, and its binding partner, FAAP20, as internal controls because both of them do not directly interact with FANCM, albeit all being present in the FA core complex^9^. In contrary to TONSL-MMS22L, no association between FANCA-FAAP20 expression and overall survival in cancer patients with wild type FANCM was observed (*P >* 0.05) (Figure 7C), while mutant FANCM-associated cancer patients with low expression of FANCA or FAAP20 exhibited poor overall survival compared to high expression version (*P <* 0.05) (Figure 7D). These data indicate that the correlation between TONSL-MMS22L low expression and favorable prognosis of cancer patients specifically depends on proficient, but not deficient FANCM; and also suggest that the FANCM-TONSL-MMS22L complex might be independent of the FA core complex in cancer prognosis.

**Figure 7.**
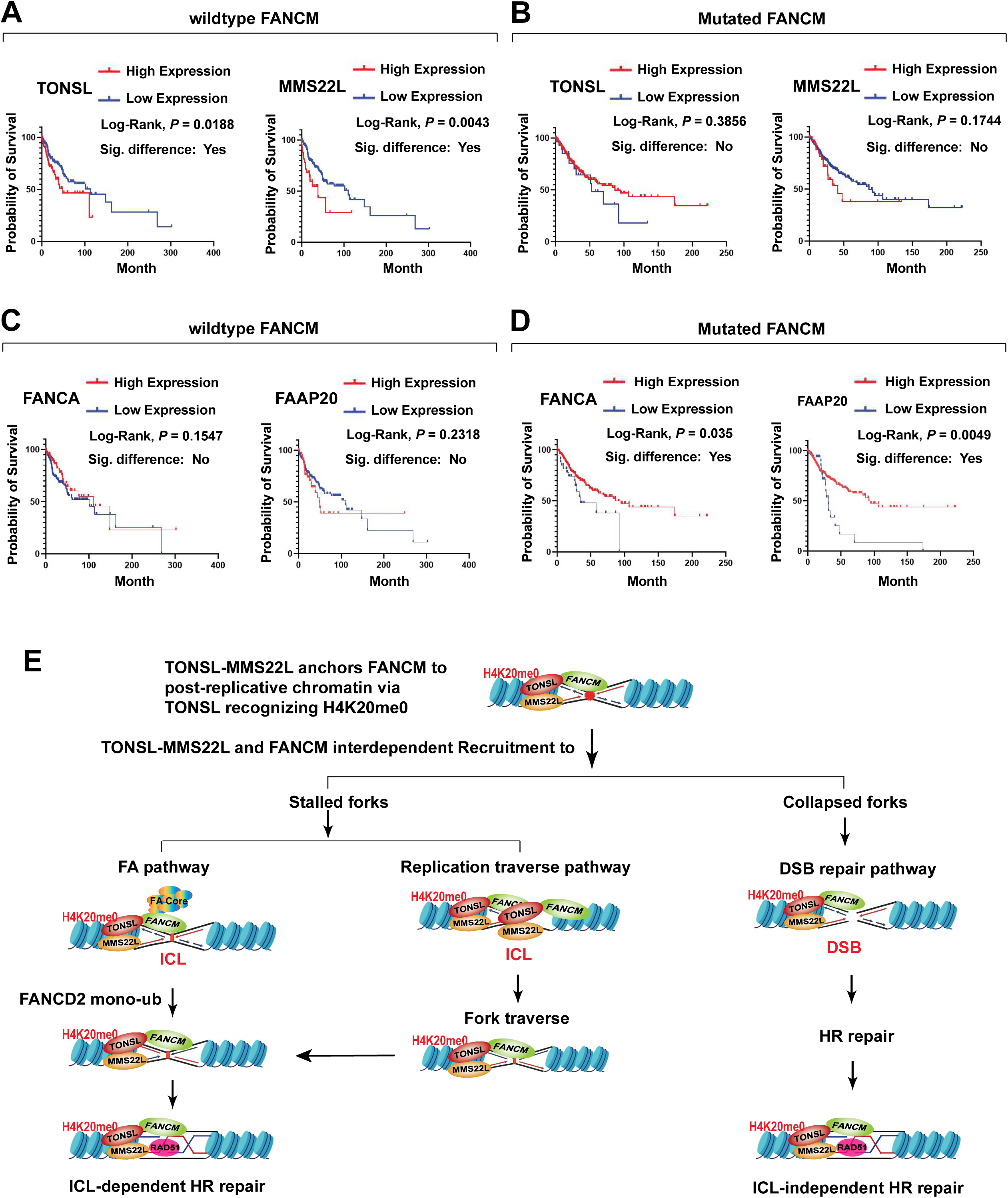
TONSL-MMS22L depends on proficient FANCM to favor prognosis of cancer patients. **(A – D)** Kaplan-Meier curves were plotted to describe the overall survival rate of patients with genes expression (TONSL, MMS22L, FANCA or FAAP20) and wildtype FANCM (A and C) or mutated FANCM (B and D). The combined 21 types of cancer from the TCGA for these survival analyses were listed, as follows: adrenocortical cancer, bladder cancer, breast cancer, cervical cancer, colon cancer, esophageal cancer, glioblastoma, head and neck cancer, kidney papillary cell carcinoma, lower grade glioma, liver cancer, lung adenocarcinoma, lung squamous cell carcinoma, ovarian cancer, pancreatic cancer, prostate cancer, melanoma, stomach cancer, testicular cancer, endometrioid cancer, uterine carcinosarcoma. **(E)** A model describes that, in response to replication stress, TONSL-MMS22L anchors FANCM to post-replicative chromatin via TONSL recognizing H4K20me0 and cooperates with FANCM to resolve replication stress by promoting activation of the FA pathway, both repair and replication traverse of ICLs, and HR repair at ICL-dependent and -independent DSBs.

## Discussion

### FANCM-TONSL-MMS22L is a chromatin-bound complex induced by replication stress

Under normal replication, we have extensively immunopurified the FANCM complex from nuclear extract of human cells and found neither TONSL nor MMS22L in the complex^12,13^. Indeed, none of other published articles reports the presence of these two proteins in the FANCM complex isolated from either nuclear extract or whole cell lysates^26,54^. Likewise, none of published data show the presence of FANCM and other FANC proteins in the TONSL-MMS22L complex purified from whole cell lysate of human cells by several laboratories^32,34,35^. This suggest that, without replication stress, the association between TONSL-MMS22L and FANCM in both nuclear extract and whole cell lysates might be weak or transient and thus not strong enough to survive from necessary washing in the IP experiments. The other possibility is that the extracted amounts of FANCM and/or TONSL-MMS22L in nuclear extract and whole cell lysates were too low to sustain a stable complex that can be immunoprecipitated for mass spectrometry analysis, because of FANCM retaining exclusively on chromatin and TONSL-MMS22L binding to histones H3-H4 incorporated into nucleosomes. In this study, we developed a chromatin-digestion followed by size exclusion chromatography protocol and treated cells with a drug that induces replication stress. By this way, we can extract soluble chromatin-bound proteins from native chromatin of human cells and concentrate endogenous FANCM complex enough for an IP-based mass spectrometry analysis. Consequently, we uncover the TONSL-MMS22L complex as a novel FANCM cofactor on chromatin. The FANCM-TONSL-MMS22L complex is constitutive because their interaction can be detected by both IP-western blotting and PLAs under normal replication, albeit weak; while sharply stimulated when cells are under replication stress. This is in agreement with the note that TONSL-MMS22L is present at normal replication forks but enhanced when forks are broken^36^.

### FANCM-TONSL-MMS22L functions on post-replicative chromatin via TONSL-H4K20me0 pathway

Recent studies have shown that, during DNA replication, TONSL-MMSL is recruited to nascent chromatin to promote repair of stalled or collapsed forks by two distinct histone-reader-dependent pathways. One is the H3.1-reader pathway where H3.1 variant binds TONSL via the TPR of TONSL and thus recruits TONSL-MMS22L to newly replicated chromatin^49,50^. The other is the unmethylated H4 at K20 (H4K20me0)-reader pathway in which the ARD of TONSL recognizes H4K20me0 in post-replicative chromatin, thereby recruiting TONSL-MMS22L to initiate HR repair by promoting RAD51^38^. Which pathway does the FANCM-TONSL-MMS22L complex choose in the cellular response to replication stress? We propose that TONSL-MMS22L may anchor FANCM to post-replicative chromatin via TONSL-H4K20me0 pathway, where TONSL-MMS22L and FANCM constitute a complex required for the cellular response to replication stress. In line with this, a mutation that inactivates TONSL binding to H4K20me0 disrupts FANCM retention on chromatin and recruitment to perturbed forks, consequently compromising cellular resistance to replication stress and attenuating the capacity of FANCM-mediated multiple DNA repair and replication traverse (Figure 7E). However, it is still possible that TONSL binding to H3.1 variant simultaneously regulates FANCM recruitment to replicated chromatin upon replication stress, which would be of interest to be investigated.

### Beyond HR at DSBs, TONSL-MMS22L resembles FANCM in resolving ICL-induced replication stress

A number of previous studies have shown that TONSL-MMS22L acts as an essential mediator to initiate HR repair by promoting RAD51 assembly on the ssDNA produced by end resection of DSBs that are generated by CPT-incurred fork collapsing or by DSB-inducing agents^32-34,36,53^. Thus far, only one paper reported that both TONSL- and MMS22L-knockdown cells displayed modest sensitivity to higher concentrations of MMC^33^, suggesting that TONSL-MMS22L might have a mild role in ICL repair. However, by creating hypomorphic mutant knockouts through CRISPR editing, we found that TONSL or MMS22L-deficient human cells exhibited marked defects in ICL repair, including hypersensitivity to MMC and cisplatin, and elevated MMC-induced chromosomal breaks and radial chromosomes, unveiling that TONSL-MMS22L resembles FANC proteins and other FA-associated proteins to play essential roles in the cellular response to ICLs (Figure 7E). This is also supported by our following lines of evidence. First, the biochemical purification reveals that TONSL-MMS22L is a FANCM cofactor but also an integral component of the FA core complex, implying that TONSL-MMS22L is an upstream actor of the FA pathway that is critical for ICL repair. Second, TONSL-MMS22L is required for normal activation of the FA pathway by promoting recruitment of the FA core complex to ICL-stalled forks and facilitating monoubiquitination and focus formation of FANCD2 in response to ICLs induced by MMC. Third, TONSL-MMS22L, like FANCM acting independently of the FA core complex, promotes replication traverse of ICLs, by which the ICL-stalled forks are not undergone collapsing and thus the replication is allowed to be continued. Therefore, beyond the established function in HR repair at DSBs^32-35^, our study expands an essentiality of TONSL-MMS22L for the cellular response to replication stress, at least, by two ways. One is working with FANCM to promote activation of the FA pathway for efficient ICL removal. The other is acting with FANCM to promote replication apparatus past over ICLs once replication forks encounter with the ICLs.

### FANCM functions in HR repair at both ICL-dependent and -independent DSBs by recruiting TONSL-MMS22L and RAD51 to perturbed replication forks

FANCM is a multifunctional DNA translocase that can dismantle the branchpoint DNA intermediates produced during DNA repair to facilitate fork reversal^3,4^, implying that FANCM might function in HR. This is supported by the facts that the FANCM ortholog Fml1 in *S. pombe* promotes gene conversion at stalled replication forks^55^, and that the FANCM ortholog Mph1in *S. cerevisiae* promotes a short homology end-directed gene conversion during DSB repair^56^. In human cells, several lines of evidence indicate that FANCM plays an important role in HR during DSB repair. First, as an upstream actor of the FA core complex, FANCM recruits the FA core complex to ICLs through its binding to ICLs and thus promotes downstream HR repair of DSBs for completion of the ICL resolution by the FA pathway^11^, suggesting that FANCM might direct ICL-dependent HR repair when replication forks stalling by the ICLs. Second, a recent study reported that FANCM along with FANCL and UBE2T were identified as a HR-promoting factor at ICL-independent DSBs by using a targeted CRISPR screen in HEK293T cells carrying the DSB-Spectrum_V2 reporter^20^. Third, our studies show that deficiency of FANCM impairs HR repair at DSBs, as detected in the DSB repair assay with U2OS cells carrying GFP-HR reporter. Forth, FANCM loss renders cells hypersensitive to CPT that drives DSBs^12,19^, suggesting that FANCM is also required for ICL-independent DSB repair, most likely, by promoting HR because FANCM and the HR mediator, TONSL-MMS22L, work in the same pathway for the cellular resistance to CPT.

Nevertheless, how might FANCM contribute to HR repair at DSBs derived from forks stalling by ICLs or from forks collapsing imposed by CPT? FANCM may work with the HR factor(s) during DSB repair. Consistent with this, FANCM forms an interdependent complex on chromatin with a HR mediator, the TONSL-MMS22L complex, which is strongly enhanced upon replication stress. FANCM recruits both TONSL-MMS22L and a key HR recombinase, RAD51 to ICL-stalled and to CPT-collapsed forks. A mutation in FANCM that inactivates its DNA translocase activity or phosphorylation disrupts such recruitments, underscoring an indispensability of FANCM for TONSL-MMS22L-RAD51 accumulation at stressed replication forks before HR repair. Importantly, FANCM and TONSL-MMS22L act together to promote RAD51 loading. Thus, FANCM recruits TONSL-MMS22L-RAD51 to promote HR repair at both ICL-dependent and -independent DSBs (Figure 7E).

### TONSL-MMS22L is a promising target for treatment of wildtype FANCM-linked cancer

Organisms have evolved a host of DNA repair systems in which numerous DNA damage response proteins cooperate to orchestrate monitoring and resolving damaged DNA^57^. Although partial or complete loss of one of these proteins impairs DNA repair, cancer cells can acquire compensatory DNA repair mechanisms for cell survival^58^. Moreover, it has been generally thought that upregulated expression of the DNA damage response proteins in cancer cells, such as RAD51^59^, renders DNA repair capacity increasing, which results in enhanced cancer progression and elevated drug resistance, consequently having poor survival of cancer patients. Conversely, downregulated expression of such proteins attenuates DNA repair capacity and thus reduced viability of cancer cells, thereby leading to a better prognosis of cancer patients. In line with this, we observed in large cohorts of cancer patients with tumors with wild type FANCM and low expression of TONSL-MMS22L have a more favorable prognosis than those with high expression. Unexpectedly, there is no correlation between TONSL-MMS22L expression and prognosis in FANCM-mutated cancer patients, suggesting that another mechanism compensates the defects of the FANCM-TONSL-MM22L complex in cancer patients carrying FANCM mutations. Thus, we propose that TONSL-MMS22L might serves as a therapeutic target for wildtype-, but not mutated FANCM-associated cancer, which needs to be further investigated.

## Method details

### Cell culture

HeLa, HEK293 and U2OS cells were grown in DMEM medium supplemented with 10% fetal bovine serum (FBS) in a 5% CO_2_ incubator at 37°C.

### Extraction of chromatin-bound protein, gel filtration, immunopurification and protein identification

Cells were treated with HU (2 mM) for 24 hours. The cell pellets were resuspended with low salt buffer (20 mM HEPES, pH 7.9, 10 mM NaCl, 3 mM MgCl_2_, 1 mM EDTA, 0.5% NP40) for 5 minutes on ice, and centrifuged at 1500 RCF for 5 min. The pellet was then resuspended with digestion buffer (20 mM HEPES, pH7.9, 100 mM NaCl, 2 mM MgCl_2_, 1mM CaCl_2_) containing microcococcal nuclease (3 U/μl and DNase I (2 U/μl) at 4°C overnight, and centrifuged at maxi speed for 15 min. All buffers were supplemented with complete protease inhibitor cocktail (Roche). The Supernatant was subjected to gel filtration on a Superose 6 column. The fractions were then harvested and immunoprecipitated with a FANCM antibody in IP buffer (20 mM HEPES, pH 7.9, 200 mM NaCl, 5 mM MgCl_2_, 0.1% Tween 20). The eluted immunoprecipitated was resolved by 10% Tris-Glycine gel, stained with silver and subjected to mass spectrometry for protein identification.

### In situ proximity ligation assay (PLA)

Cells were grown on Mattek glass bottomed plates then treated with 1.5 µM TMP/UVA or UVA only. UVA exposure was carried with a Rayonet chamber at 3 J/cm2. With fresh medium incubation for 60 min, cells were followed by treated with 0.1% formaldehyde for 5 min and twice CSK-R buffer (10 mM PIPES, pH 7.0, 100 mM NaCl, 300 mM sucrose, 3 mM MgCl2, 0.5% Triton X-100, 300 µg/ml RNAse), and fixed in 4% formaldehyde in PBS for 10 min, then followed by incubation in pre-cold methanol for 20 min at -20 °C, 100 ug/ml RNAse for 30 min at 37 °C, and then 0.5% Triton X-100 for 10 min at 4°C. In situ PLA was performed using the Duolink PLA kit (Sigma-Aldrich) according to the manufacturer’s instructions. Briefly, cells were blocked for 30 min at 37 °C and incubated with the respective primary antibodies for 30 min at 37 °C. Following three times washing with PBST (PBS plus 0.1% Tween 20), PLUS and MINUS PLA probes were coupled to the primary antibodies for 1 h at 37 °C. After three times washing with buffer A (0.01 M Tris, 0.15 M NaCl, and 0.05% Tween-20) for 5 min, PLA probes ligated for 30 min at 37 °C. were Then amplification using detection was performed at 37 °C for 100 min and washed with wash buffer B (0.2 M Tris 0.1 M NaCl) for three times. Finally, they were coated with mounting medium containing DAPI (Prolong Gold, Invitrogen). The stage of the S phase was determined by immunostaining of cells with an antibody against PCNA conjugated with Alexa 647. Then PLA foci was imaged on Nikon TE2000 spinning-disk confocal microscope.

### Generation of heterozygous knockout cell lines by CRISPR/Cas9

HeLa cells were transfected with pSpCas9(BB)-2A-Puro (PX459) V2.0 plasmid (Addgene, #62988) inserted with TONSL and MMS22L guide RNMA sequences, respectively. After 48 h, cells were selected with puromycin (2 μg/ml). Individual clones were picked up, cultured and screened by western blotting, followed by PCR of genomic DNA. The amplified DNA fragments were cloned into pGEM-T Easy vector, respectively. Sanger sequencing was then performed. TONSL gRNA targeting exon 3: GAGCGCGCTGACGACCCTCT. MMS22L gRNA targeting exon 2: CCTGACTGACAGCTTAGAGC. HeLa cells were transfected with pSpCas9(BB)-2A-Puro (PX459) V2.0 plasmid only and then selected by puromycin at the same concentration. The survived clones were used as a wildtype control, marked as HeLa WT, because both TONSL and MMS22L are wildtype.

### Plasmid construction and transfection

The expression plasmid pIRES-Flag-FANCM, pIRES-Flag-TONSL or pIRES-Flag-MMS22L used for generation of stable cell lines was constructed by inserting FANCM, TONSL or MMS22L cDNA with a FLAG tag at its N-terminus into pIRES-Neo3 (Clontech). pIRES-Flag-TONSL deletion mutants and N571A point mutant generated by PCR were constructed by inserting TONSL deletion fragment and point mutant fragment with a FLAG tag at their N-terminus into pIRES-Neo3, respectively. Lipofectamine 2000 (Thermo Fisher Scientific) or GenJet™ *In Vitro* DNA Transfection Reagent (SignaGen Laboratories) was used for transfection in HEK293 cells or HeLa cells.

### siRNA experiments

Cells were transfected with siRNA oligos using Lipofectamine 2000 (ThermoFisher Scientific) according to manufacturer’s manual. After 2 days post-transfection, cells were used for various assays. TONSL siRNA oligo #1 (CCUAAUAAAUGAAGCUGCU)^33^, TONSL siRNA oligo #2 (GAGCUGGACUUAAGCAUGA)^36^. MMS22L siRNA oligo #1 (GGUAGAAGAUGUUGCAAGU)^36^, MMS22L siRNA oligo #2 (CCCUUAAUGAUACGACGAA)^36^. siFANCA oligo (AAGGGUCAAGAGGGAAAAAUA)^60^. siFANCM oligo (AGGCUGUGCAACAAGUUAU)^8^. siControl oligo (UUCUCCGAACGU GUCACGUTT). All oligos were purchased from GenePharma (Shanghai, China).

### Cell extraction and subcellular fractionation

Cells were resuspended with the lysis buffer for the preparation of whole cell extract as described. For cellular fractionation, the cells were washed with PBS twice and were lysed in CSK (10 mM PIPES, pH 6.8, 100 mM NaCl, 300 mM sucrose, 3 mM MgCl2, 0.5% Triton X-0.5% Triton X-100) supplemented with complete protease inhibitors (Roche) for 5 min on ice. Following centrifugation at 1500 RCF for 5 min, the pellet was treated with 2X sample loading buffer (BioRad) and boiled for 5 min. After centrifugation, the supernatant was used as chromatin fraction (chromatin-bound proteins).

### Immunofluorescence (IF)

Cells were plated onto glass coverslips in 12-well plates. After one or two days, cells were treated with the appropriate drugs as indicated. Cells were pre-extracted twice with CSK (100 mM NaCl, 300 mM sucrose, 3 mM MgCl2, 10 mM PIPES, pH 6.8) supplemented with 0.5% Triton X-100 for on ice for 5 min twice and then fixed with 2% paraformaldehyde (PFA) and permeabilized with 0.5% Triton X-100 in PBS for 10 min on ice (for FANCM, FANCA and RAD51) or at room temperature (MMS22L and TONSL). After blocking with the blocking buffer (1% BSA, 5% goat serum, 0.2% Triton X-100 in PBS) for 1h at room temperature, cells were incubated with primary antibody at 37°C for 1.5 h, and then washed with PBST (0.2%Trion-X100 in PBS). Cells were blocked with blocking buffer for 1h again and then incubated with goat anti-rabbit secondary antibody (Vector Laboratories) at 37°C for 1h. The coverslips were mounted with VECTASHIELD plus Antifade Mounting Medium with DAPI (Vector Laboratories). Images were acquired using a Zeiss Axio Observer 5 microscope (Carl Zeiss). For TONSL IF, cells were fixed with pre-cooled methanol for 10 min followed by 10 min with 2% paraformaldehyde (PFA) on ice. For FANCD2 IF, cells were fixed in 2% paraformaldehyde in PBS for 10min on ice. After being washed with PBS, the cells were permeabilized with 0.5% Igepal CA-630 in PBS for 10min and washed again with PBS. After blocking with 1% BSA, 5%Goat in PBS for 1h at room temperature, cells were incubated with primary antibodies at 37°C for 1.5h and secondary antibodies at 37°C for 1h. The coverslips were then washed with PBST (0.05%Tween 20 in PBS).

### Clonogenic survival assay

The clonogenic survival assays were performed as described ^8^ with a modification. Briefly, after 16-18 h of seeding, cells were exposed to MMC (Sigma) or CPT (MedChemExpress) at the indicated concentrations for 10-12 days. The clones were fixed with methanol/acetic acid (10:1), stained with 0.1% crystal violet and counted.

### Chromosome breakage assay

Cells were plated onto 60 mm dishes for one day and then treated with MMC (40 ng/ml) for 24h, before being incubated with colcemid (0.1μg/ml) for 2.5 h. Cells were harvested and resuspended with 0.56% KCl at 37°C for 25 min and fixed with methanol/acetic acid (3:1). Metaphase spreads were prepared on glass slides, immersed in Gurr’s buffer (PH6.8) containing Giemsa solution (Sigma) (9:1) for 18min. The slides were rinsed with water and mounted with Eukitt Quick-hardening mounting medium (Sigma).

### SCE assay

Cells were plated on to 60 mm dishes for one day and then treated with 100 μM BrdU for two cell cycles, before being incubated with colcemid (0.1 μg/ml) for 3 h. Cells were harvested, resuspended with KCl and fixed as described in the chromosome breakage assay. Metaphase spreads were prepared on glass slides and then incubated in Gurr’s buffer (PH6.8) containing Hoechst33258 for 30 min. The slides were rinsed with Macllvaine solution (PH7.0) and then covered with coverslips. The slides were irradiated with black light for 1 h and then washed with 2x SSC (PH7.0) at 62°C for 1 h. The slides were stained with Giemsa and sealed as described in the chromosome breakage assay.

### Traverse assay

Cells were incubated for 25 min with 5 μM Dig-TMP before UVA irradiation in a Rayonet chamber at 3 J/cm^2^ for 5 min. Cells were then incubated in 20 μM CldU for 20 min, followed by 200 μM IdU for 20 min. After the replication labeling, Cells were harvested and mixed with lysis buffer (0.5% SDS in 200 mM Tris-HCl [pH 7.5], 50 mM EDTA) on a glass slide. After DNA spreading, the slides were air-dried, fixed in 3:1 methanol/acetic acid, incubated in 2.5 M HCl for 60 min, neutralized in 0.4 M Tris-HCl (pH 7.5) for 10 min, washed in PBST (0.05% Tween 20 in PBS), and immunostained. incubated in 2.5 M HCl for 60 min, neutralized in 0.4 M Tris-HCl (pH 7.5) for 5 min, washed in PBS, and immunostained with antibodies at room temperature as following: rat anti-BrdU (CldU), 1:200; Dylight 649 goat anti-rat, 1:100; mouse anti-BrdU (IdU), 1:40 and chicken anti-digoxigenin, 1:200; and Dylight 488 goat anti-mouse, 1:100 and Qdot 655 goat anti-chicken, 1:2,000. Imaging was acquired using a Zeiss Axiovert 200 M microscope with the Axio Vision software. The quantum dot signals were imaged with a Qdot 655 filter.

### GFP-based HR repair assay

U2OS cells carrying GFP-based HR reporter construct were kindly provided by Dr. Jeremy Stark at the City of Hope and seeded in 6 well plates. On the next day, cells were transfected with FANCM siRNA, TONSL siRNA, MMS22L siRNA and control siRNA as indicated. After one day, cells were transfected with the I-SceI plasmid (Addgene) in the presence with siRNA, using lipofectamine 2000. After 2 days, cells were harvested by trypsinization and analyzed by flow cytometry.

### Data acquisition and analysis of TCGA cancer patients

The clinical, mRNA expression Z-scores (RNASeq V2 RSEM) and genomic alteration data of TCGA Pan-Cancer study for 32 cancer types were downloaded from the cBioPortal for Cancer Genomics (https://www.cbioportal.org)^61^. Patients were stratified into low- and high-expression groups for TONSL, MMS22L, FANCA or FAAP20, based on optimal expression cut-off values derived from survival outcomes using the X-tile tool. The survival analyses of TCGA patients with wildtype or mutated FANCM were performed. Kaplan-Meier survival curves were plotted using GraphPad Prism. The log-rank test was used to calculate the significance of survival-time differences between two group of patients with a threshold of *P*-value < 0.05.

## Acknowledgement

We thank Dr. Daniel Durocher at Samuel Lunenfeld Research Institute, Canada for plasmids carrying TONSL and MMS22L cDNAs, and Dr. Jeremy Stark at the City of Hope Medical Center, USA for U2OS cell line carrying HR reporter construct. This work was supported by the National Natural Science Foundation of China (32070716 to Z.Y.), in part by the Oncology Discipline Group, the Second Affiliated Hospital and Yuying Children’s Hospital, Wenzhou Medical University, China (to H. Z), and in part by the Intramural Research Program of the National Institute on Aging (AG000688[2024] to W.W., and Z01-AG000746-08 to M.M.S), National Institutes of Health, USA.

## Materials Availability

Further information and requests for reagents and resources should be directed to and will be fulfilled by the lead contact. Zhijiang Yan

## Supplemental Information

**Figure S1 (Related to Figure 1).**
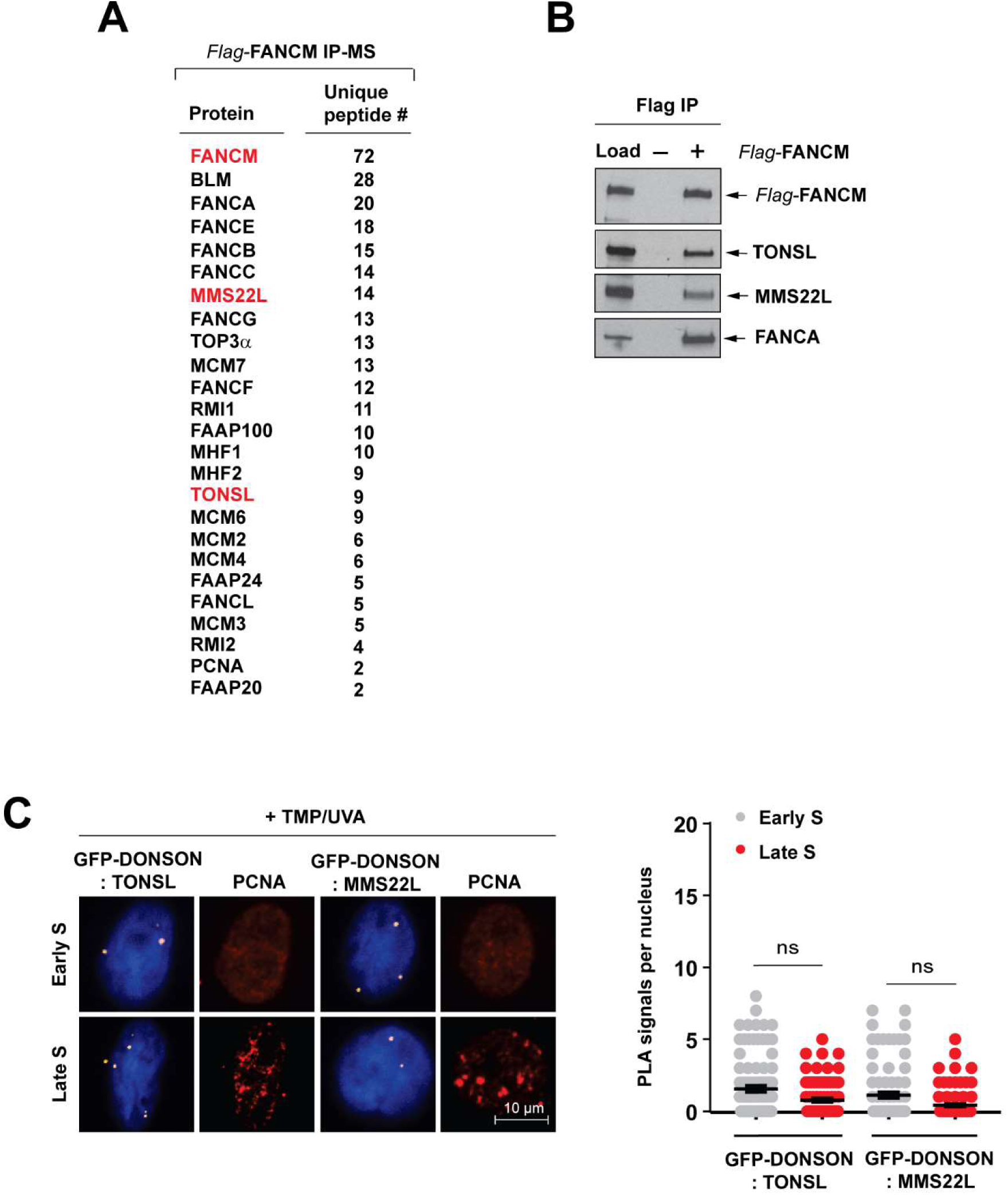
Replication stress stimulates the FANCM-TONSL-MMS22L complex on chromatin. **(A)** We extracted soluble chromatin-bound proteins from native chromatin of HEK293 cells stably expressing Flag-tagged FANCM by treatment with benzonase, a pan nuclease degrading all types of DNA and RNA, and then purified the FANCM complex with a Flag antibody. Mass spectrometry reveals the proteins identified in the FANCM precipitate and the number of unique peptides derived from the corresponding proteins. **(B)** Immunoblotting shows the presences of FANCM, TONSL, MMS22L and FANCA in the precipitate described in (A). HEK293 cells without expression of Flag-FANCM were used as a control. **(C)** *Left panel*: Representative images of the PLA between GFP-DOSON and TONSL or MMS22L in early and late phase cells treated with TMP/UVA. *Right panel*: A graph showing the frequencies of PLA signals described in (C, *Left panel*). Data are mean ± SEM from three biological replicates. Number of nuclei: PLA between GFP-DONSON and TONSL, early S phase cells = 92, late S phase cells = 84; PLA between GFP-DONSON and MMS22L, early S = 86, late S = 87.

**Figure S2 (Related to Figure 2).**
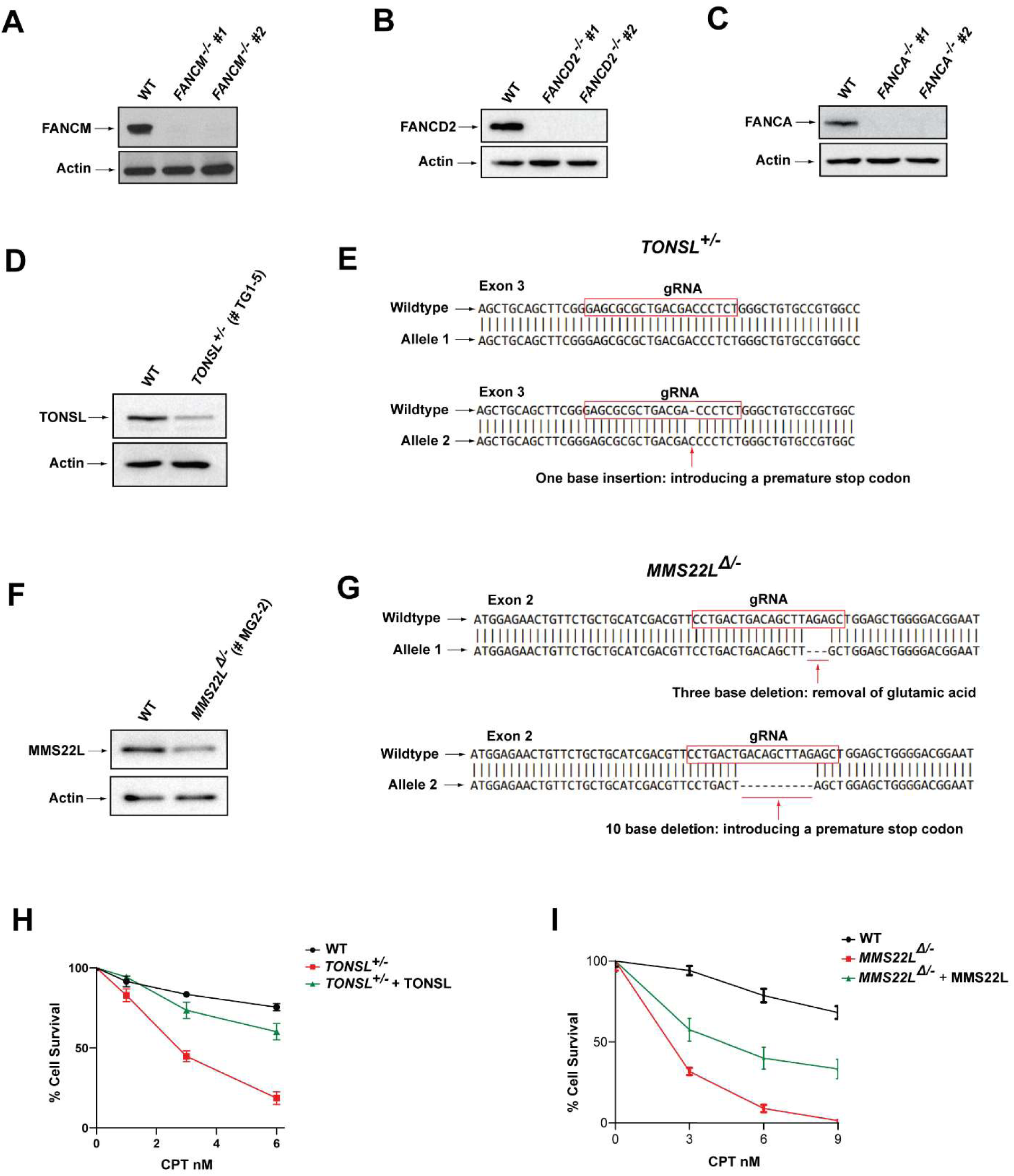
Both TONSL and MMS22L heterozygotes are hypersensitive to replication stress. **(A)** Immunoblotting shows the level of FANCM protein in HeLa wildtype (WT) cells and HeLa cells-derived clones by CRISPR/Cas 9 genome editing using two different gRNA oligos to FANCM locus. Actin was used as a loading control. **(B and C)** As described in (A), except FANCD2 (B) and FANCA (C). **(D and F)** As described in (A), except one gRNA oligo to TONSL locus (D) and one gRNA to MMS22L locus (F). **(E)** Sanger sequencing of the clone TG1-5 showing that one allele is the same as wild type, while the other carries a deletion of one nucleotide within the gRNA-binding sequence that introduces a premature stop codon. These results establish that TG1-5 is a heterozygous knockout clone (*TONSL^+/-^*). **(G)** As described in (E), except that one allele in the clone MG2-2 carries a deletion of three nucleotides within gRNA-binding sequence, resulting in removal of a glutamic acid; while the other carries a deletion of 10 nucleotides that introduces a premature stop codon. Therefore, MG2-2 is a compound heterozygous knockout clone (*MMS22L*^Δ/-^). **(H)** Clonogenic survival assays of HeLa WT cells, *TONSL^+/-^* cells and *TONSL^+/-^* cells complemented with wildtype TONSL following CPT treatment at the indicated concentrations. Data are mean ± SEM from three independent experiments. Each experiment has triplicate cultures. **(I)** As described in (H), except MMS22L as indicated.

**Figure S3 (Related to Figure 2).**
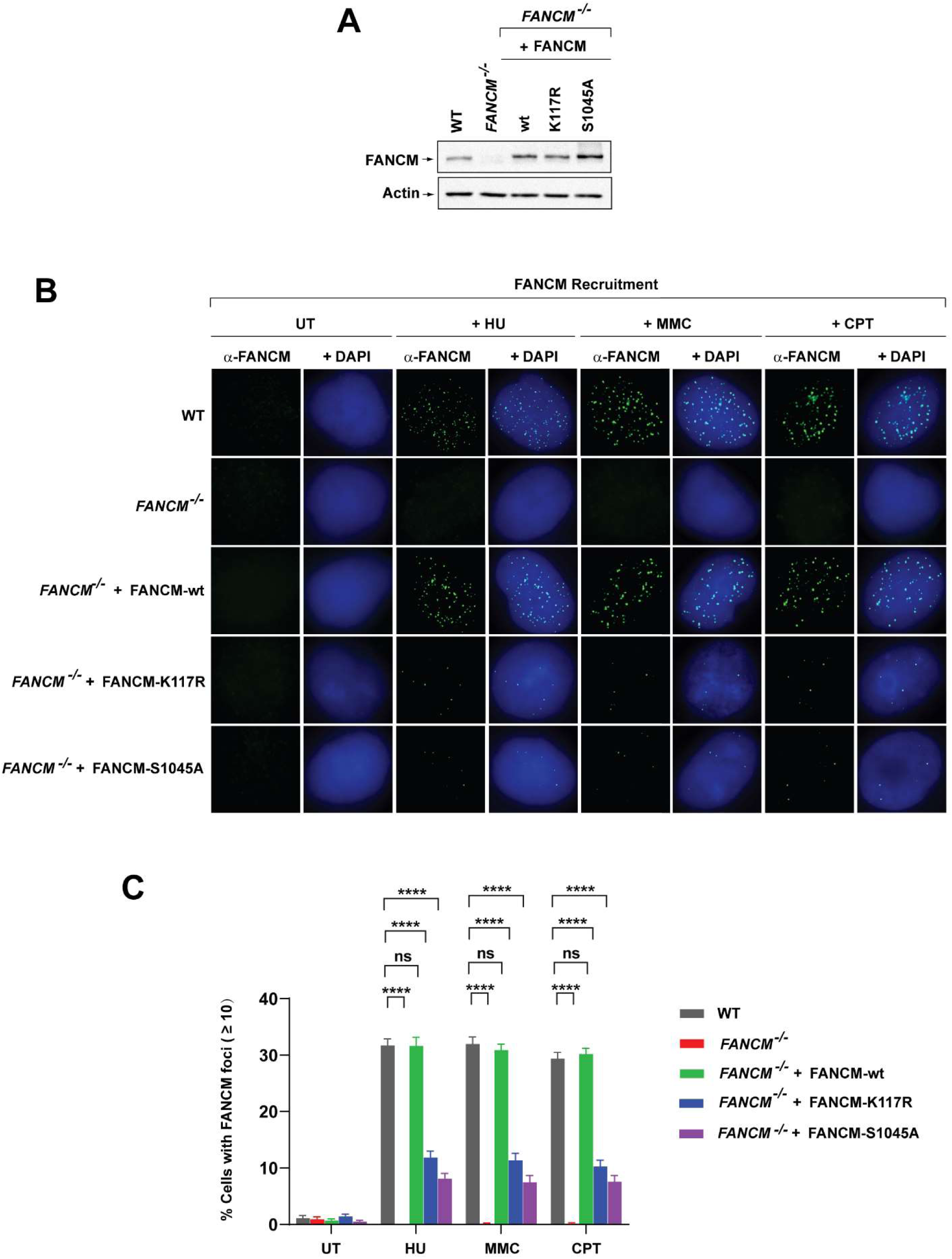
Both DNA translocase activity and phosphorylation of FANCM are required for its efficient recruitment to stalled and collapsed forks. **(A)** Immunoblotting shows the level of FANCM protein in HeLa wildtype (WT) cells, *FANCM^-/-^* cells and complemented versions with FANCM wildtype (wt), DNA translocase-inactivated mutant FANCM-K117R or phosphorylation-abolished mutant FANCM-S1045A. Actin is an internal control. **(B)** Representative immunofluorescence images showing FANCM nuclear foci in various cell types as described in (A). Cells were treated with HU (2 mM) for 24 h, MMC (60 ng/ml) for 18 h, and CPT (1.5 μM) for 24 h. **(C)** A statistical graph showing the mean values of the percentage of FANCM-foci-positive cells in untreated (UT) and drug-treated cells with standard errors of the mean (SEM) from three independent experiments. A cell containing more than ten foci was considered as foci-positive. At least 200 nuclei were counted for each cell line. Approximate 30% of WT cells revealed more than 10 of large and discrete nuclear foci containing FANCM after exposure to HU, MMC or CPT. In contrast, little few FANCM foci were detected in control cells without drug treatment or in HeLa-derived FANCM null (*FANCM^-/-^*) cells treated with the same drugs. The absence of FANCM foci in *FANCM^-/-^* cells were all complemented by reintroduction of wildtype FANCM (WT) to the levels equivalent to those in WT cells. Of note, re-expression of FANCM-K117R or FANCM-S1045A restored only ∼ 11% and 7.5 % of *FANCM ^-/-^* cells with more than 10 of FANCM foci, respectively, indicating that both DNA translocase activity and phosphorylation of FANCM are required for to its efficient recruitment to stalled and broken forks in human cells.

**Figure S4 (Related to Figure 2).**
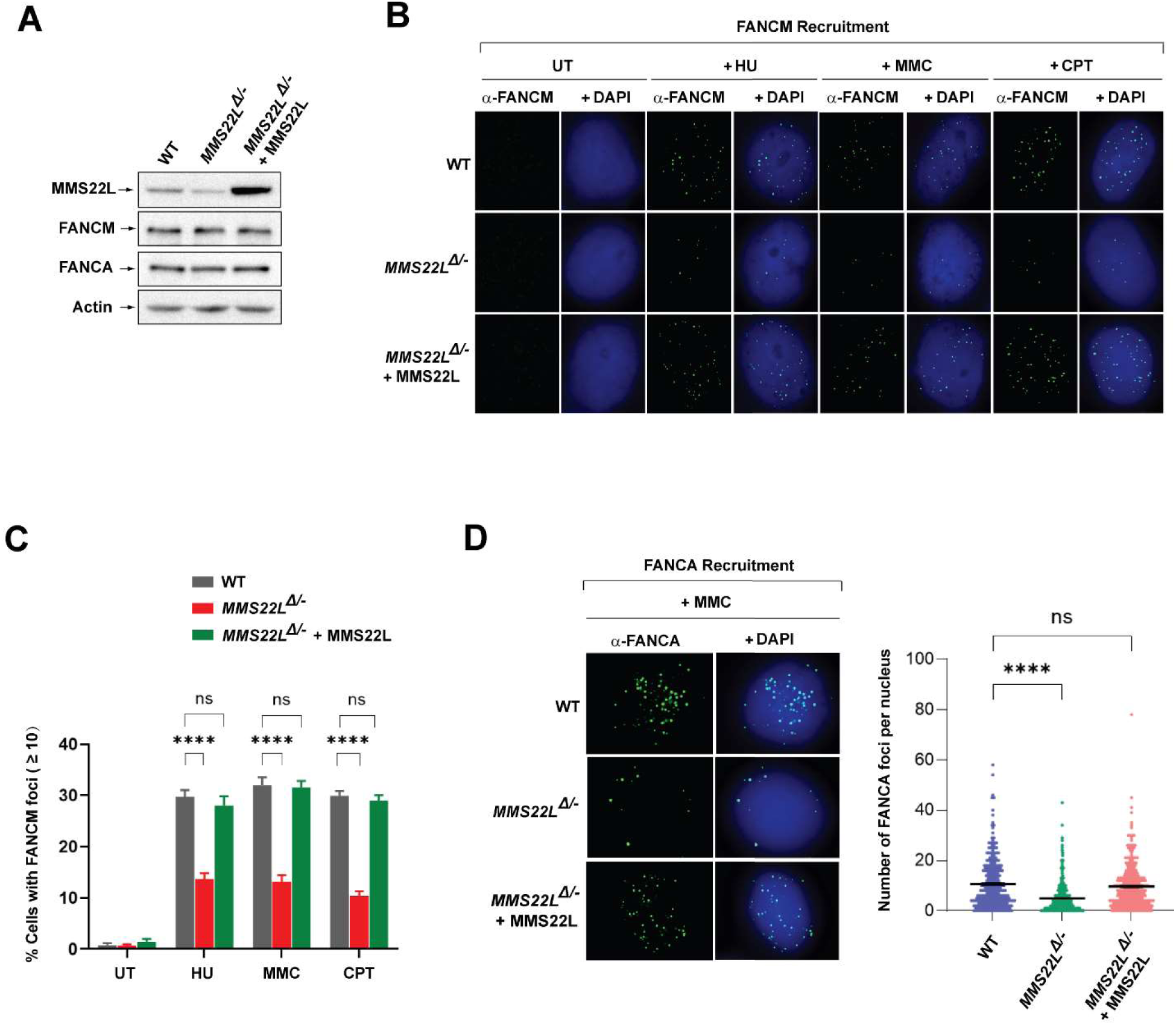
MMS22L promotes recruitment of FANCM and the FA core complex to stalled and collapsed forks. **(A)** Immunoblotting shows the levels of MMS22L, FANCM, FANCA in whole cell lysate from wild-type (WT) HeLa cells, *MMS22L*^Δ/-^, and *MMS22L*^Δ/-^ cells complemented with MMS22L. Actin was included as a loading control. **(B)** Immunofluorescence images showing FANCM nuclear foci in various cells described in (A) after treatment with HU (2 mM) for 24 h, MMC (60 ng/ml) for 18 h or CPT (1.5 μM) for 24 hr. **(C)** A statistical graph shows the mean values of the percentage of FANCM-foci-positive cells in untreated (UT) and drug-treated cells from three independent experiments with standard errors of the mean (SEM). A cell containing more than ten foci was considered as foci-positive. At least 200 nuclei were counted for each cell line. The error bars are standard error of the mean (SEM) from three independent experiments. **** represents *P* < 0.0001. “ns” represents non-significant difference. **(D)** As described in (B and C), except a FANCA antibody and MMC were used.

**Figure S5 (Related to Figure 3).**
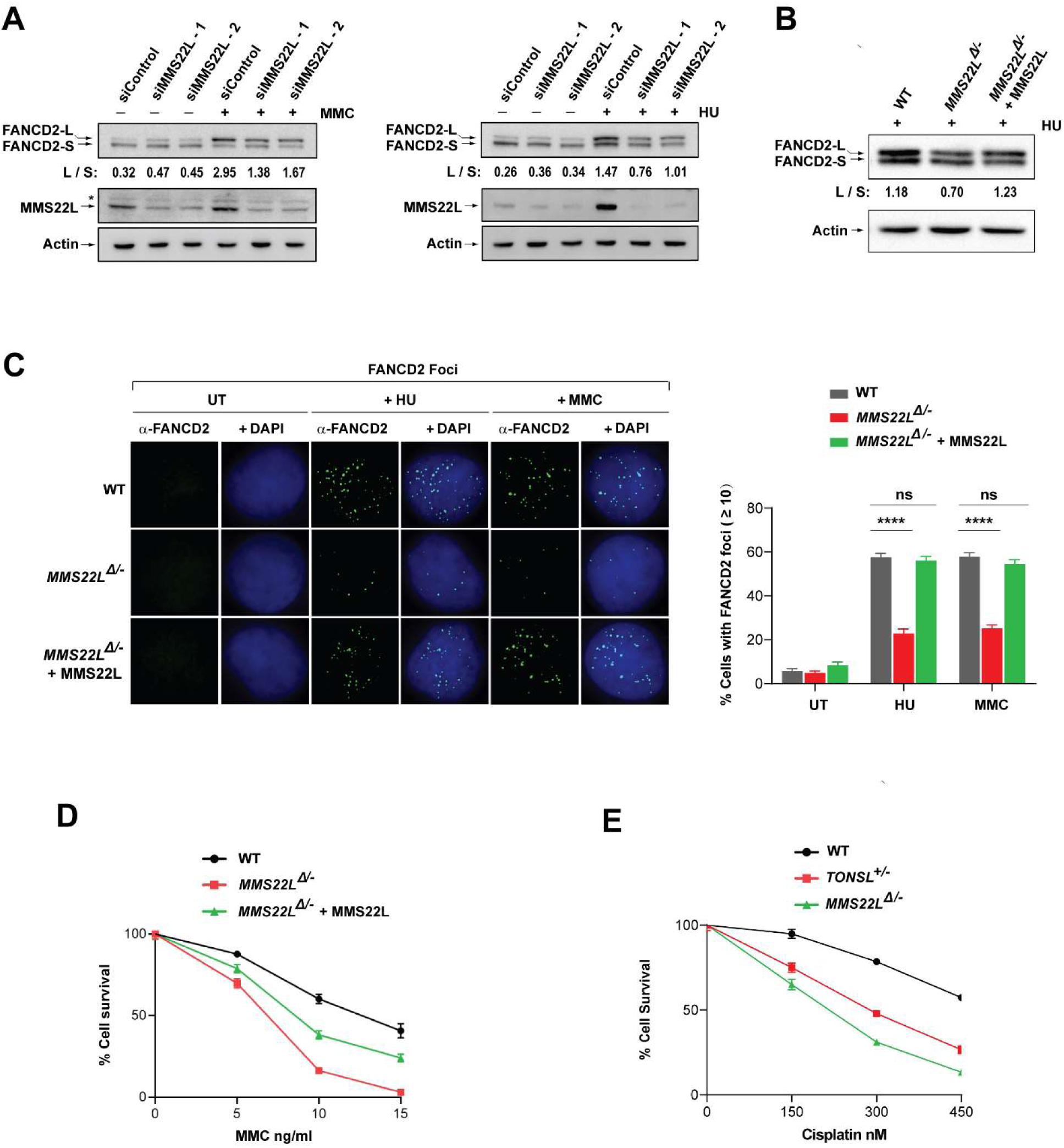
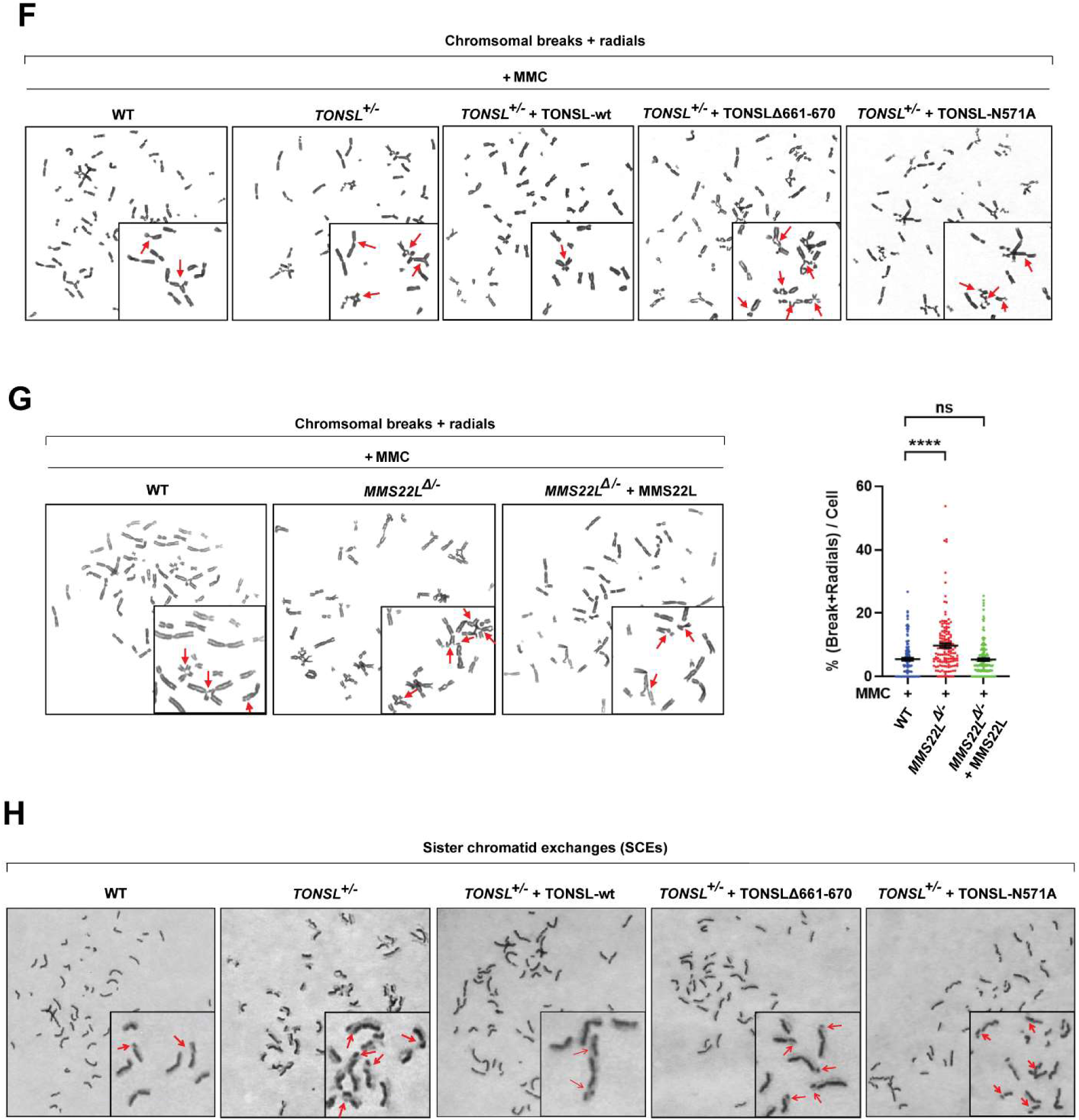
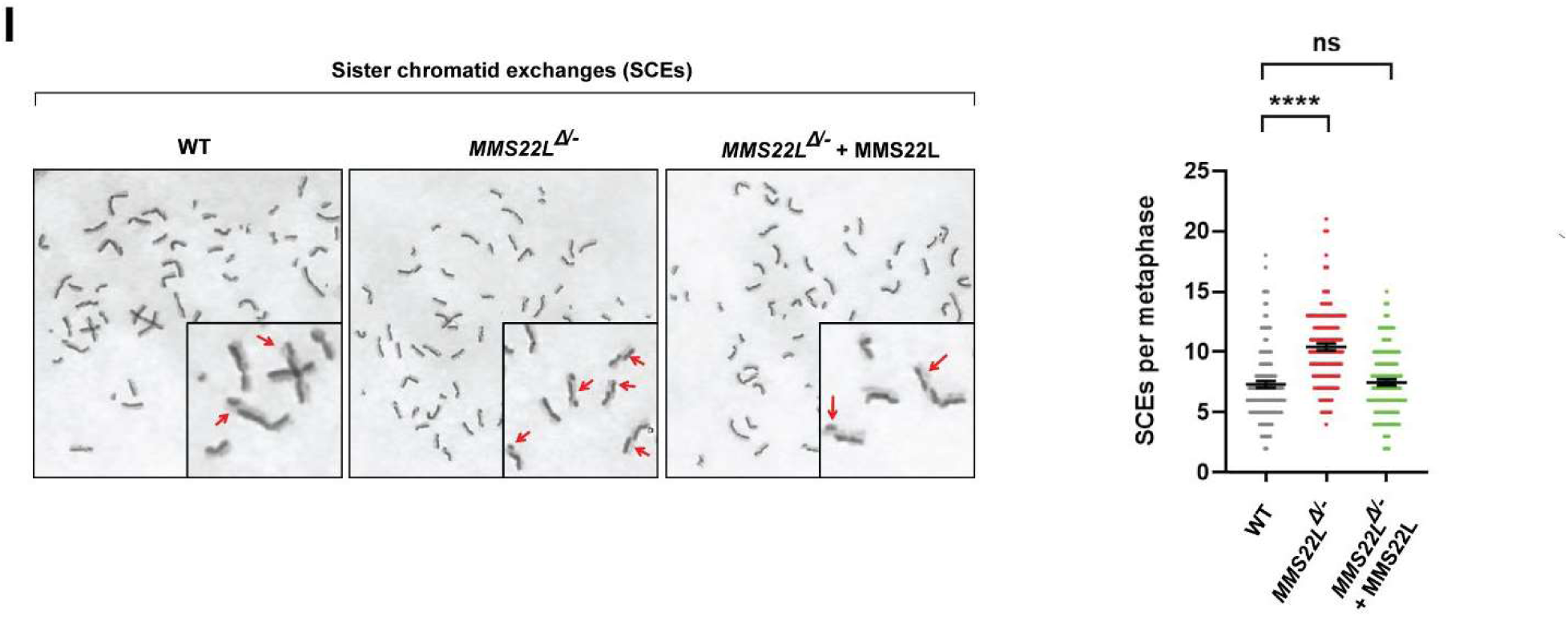
TONSL-MMS22L complex promotes activation of the FA pathway, ICL repair and SCE suppression. **(A)** Immunoblotting shows that HeLa cells depleted of MMS22L have a reduced level of monoubiquitinated FANCD2 in the presence of MMC (60 ng/ml) for 16 h (*Left panel*) or HU (2 mM) for 16 h (*Right panel*). “L” (long) and “S” (short) represent ubiquitinated and non-ubiquitinated forms, respectively. The ratio between long and short forms was obtained using Image J Software and shown below the blots. Actin was used as a loading control. **(B)** As described in (A), except HeLa wildtype cells, *MMS22L*^Δ/-^ cells and *MMS22L*^Δ/-^cells complemented with wildtype MMS22L. **(C)** Immunofluorescence images (*Left panel*) and a quantification graph (*Right panel*) showing FANCD2 nuclear foci in various cells described in (B) after treatment with HU (2 mM) for 16 h or MMC (60 ng/ml) for 16 h. Data are mean ± SEM from three independent experiments. **** represents *P* < 0.0001. “ns” represents nonsignificant difference. **(D** and **E)** Clonogenic survival assays of indicated cells following treatment with MMC or cisplatin at the indicated concentrations. **(F)** Representative images showing chromosomal breaks and radial chromosomes in MMC treated various cells as indicated. **(G)** *Left panels:* As described in (F*). Right panels:* A graph showing the number of MMC-induced chromosomal breaks and radial chromosomes in each metaphase cell as indicated. Data are mean ± SEM from three independent experiments. About 50 metaphase cells were counted per each sample in each experiment. **** represents *P* < 0.0001. “ns” represents nonsignificant difference. **(H)** Representative images showing the number of sister chromatid exchanges in each metaphase cells as indicated. **(I)** *Left panels:* As described in (H). *Right panels:* A graph showing the levels of SCEs in various cells as indicated. Data are mean ± SEM from three independent experiments. About 50 metaphase cells were counted per each sample in each experiment. **** represents *P* < 0.0001. “ns” represents nonsignificant difference.

**Figure S6 (Related to Figure 4).**
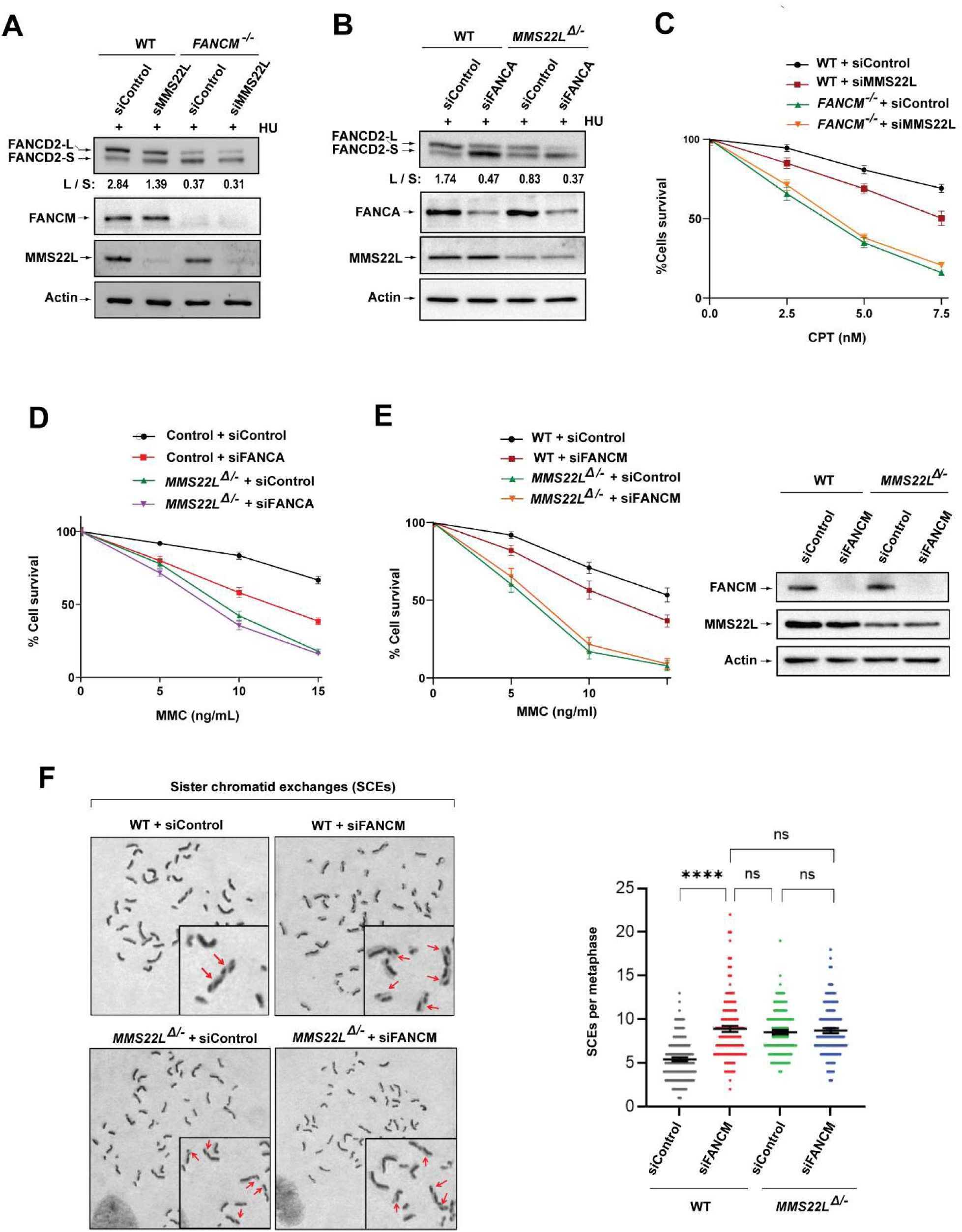
TONSL-MMS22L works together with FANCM and the FA core complex for FANCD2 monoubiquitination, cellular resistance to replication stress and SCE suppression. **(A)** Immunoblotting shows levels of monoubiquitinated and unubiquitinated FANCD2, FANCM, MMS22L in whole cell lysates from various cells as indicated on the top. Cells were treated with HU (2 mM) for 18 h. Actin was included as a loading control. **(B)** As described in (A), excepted FANCA. **(C)** Clonogenic survival assays of various cells as indicated following CPT treatment at the indicated concentrations. Data are mean ± SEM from three independent experiments. Each experiment has triplicate cultures. **(D)** As described in (C), except that MMC was used. **(E)** *Left panels:* As described in (C), except MMC was used. *Right panels:* Immunoblotting shows the levels of FANCM and MMS22L in whole cell lysates from various cells as indicated. Actin was used as a loading control. **(F)** *Left panel:* Representative images showing SCEs (red arrows) in various cells as indicated. *Right panel:* A graph showing the spontaneous SCE levels of various cells as indicated. Data are mean ± SEM from three independent experiments. About 50 metaphase cells were counted per each sample in each experiment. **** represents *P* < 0.0001. “ns” represents nonsignificant difference.

**Figure S7 (Related to Figure 6).**
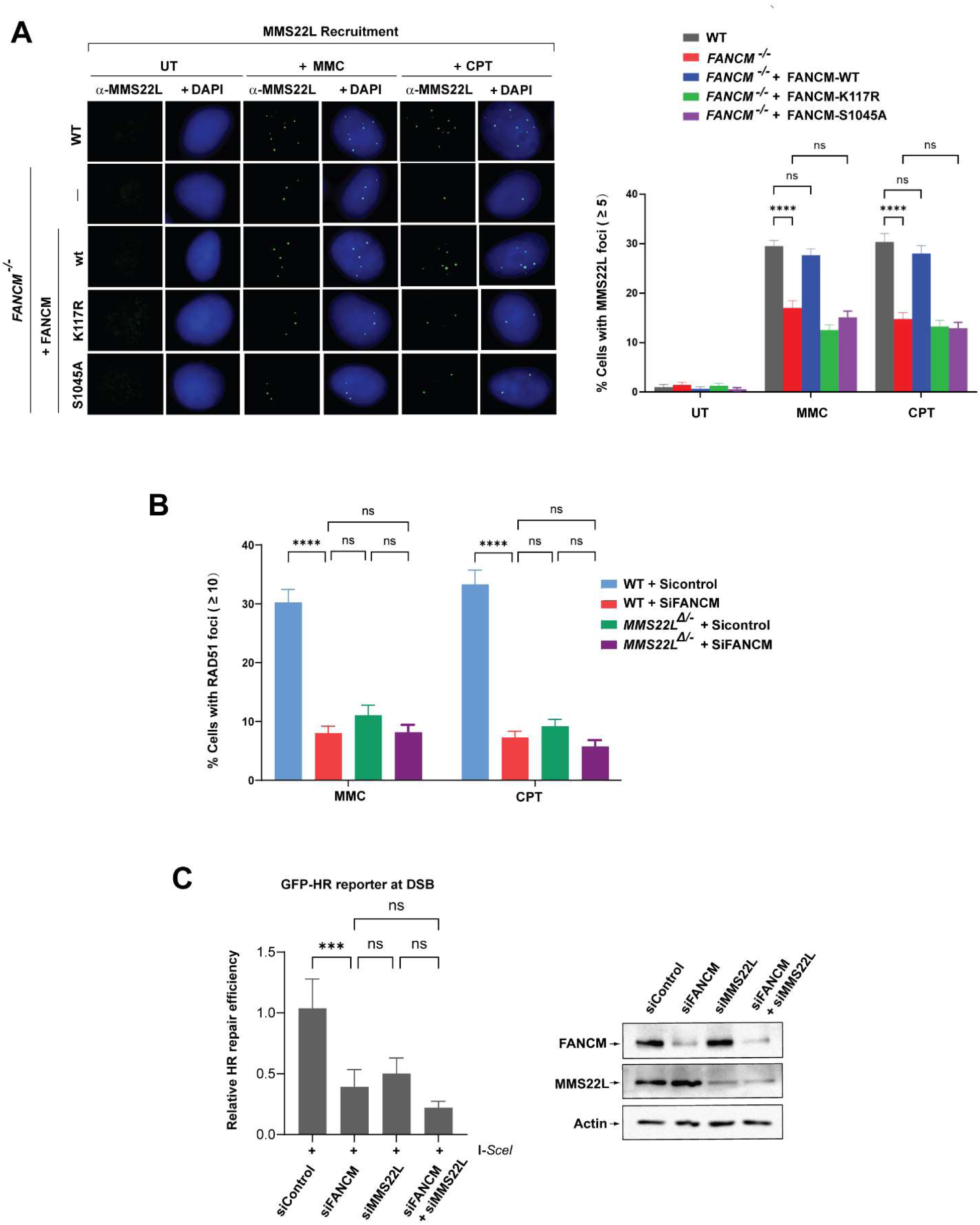
FANCM and TONSL-MMS22L function together to promote homologous recombination (HR) repair. **(A)** Representative images showing MMS22L nuclear foci Immunofluorescence images (*Left panels*) and a quantification graph (*Right panels*) showing MMS22L nuclear foci in various cells after treatment with MMC (60 ng/ml) for 18 h or CPT (1.5 μM) for 24 h. Data are means ± SEM from three independent experiments. **** represents *P* < 0.0001. “ns” represents nonsignificant difference. **(B)** As described in (A), except RAD51 nuclear foci. **(C)** *Left panels:* A quantitation graph showing relative repair efficiency derived from the positive GFP cells in U2OS-DR-GFP-reporter cells transfected with various siRNA oligos as indicated. Data are the mean with SEM from three independent experiments. *** represents *P* < 0.001. “ns” represents nonsignificant difference. *Right panels:* Immunoblotting shows the level of FANCM and MMS22L in whole cell lysates of various cells as indicated. Actin was included as a loading control.

